# Transcriptional heterogeneity of ventricular zone cells throughout the embryonic mouse forebrain

**DOI:** 10.1101/2021.07.05.451224

**Authors:** Dongjin R. Lee, Christopher T. Rhodes, Apratim Mitra, Yajun Zhang, Dragan Maric, Ryan K. Dale, Timothy J. Petros

## Abstract

The ventricular zone (VZ) of the nervous system contains radial glia cells that were originally considered relatively homogenous in their gene expression. However, a detailed characterization of transcriptional diversity in these VZ cells has not been reported. Here, we performed single-cell RNA sequencing to characterize transcriptional heterogeneity of neural progenitors within the VZ and subventricular zone (SVZ) of the mouse embryonic cortex and ganglionic eminences (GEs). By using a transgenic mouse line to enrich for VZ cells, we detect significant transcriptional heterogeneity within VZ and SVZ progenitors, both between forebrain regions and within spatial subdomains of specific GEs. Additionally, we observe differential gene expression between E12.5 and E14.5 VZ cells, which could provide insights into temporal changes in cell fate. Together, our results reveal a previously unknown spatial and temporal genetic diversity of telencephalic VZ cells that will aid our understanding of initial fate decisions in the forebrain.

## INTRODUCTION

During embryogenesis, the telencephalon is derived from a neuroepithelium sheet at the anterior end of the neural plate. Along the neuraxis, the telencephalon develops an anterior-posterior and dorsal-ventral identity guided by extrinsic signals (Wilson and Houart, 2004). The dorsal telencephalon gives rise to excitatory glutamatergic cells of the cortex (CTX) whereas the ventral telencephalon is comprised of three transient structures, the lateral, medial and caudal ganglionic eminences (LGE, MGE & CGE, respectively), that give rise to all forebrain GABAergic cells (Hebert and Fishell, 2008). Each of these regions contains progenitor domains known as the ventricular zone (VZ), which harbors *Nestin*-expressing radial glia cells and apical progenitors (APs), and the subventricular zone (SVZ) where basal progenitors (BPs) reside (Batista-Brito et al., 2008; Flames et al., 2007; Inan et al., 2012; Long et al., 2009; Petros et al., 2015; Rowitch and Kriegstein, 2010; Turrero Garcia and Harwell, 2017; Wonders et al., 2008). Early studies supported the hypothesis that common RGC progenitors in the cortex become progressively restricted over future cell divisions (Desai and McConnell, 2000; Gao et al., 2014; McConnell and Kaznowski, 1991). More recent experiments identified some heterogeneity of forebrain VZ cells (Franco and Muller, 2013; Llorca et al., 2019), but a comprehensive characterization of these VZ neural progenitor cells has not been performed.

There is a significant expansion in the number of SVZ cells as embryogenesis progresses which is likely related to developmental changes in cell fate (Lui et al., 2011). Several mechanisms have been shown to guide cell fate decisions in the ventral telencephalon, including spatial gradients of signaling factors (Brandao and Romcy-Pereira, 2015; Flames *et al*., 2007; McKinsey et al., 2013; Tyson et al., 2015; Wonders *et al*., 2008; Xu et al., 2010), cellular birthdates (Bandler et al., 2017; Inan *et al*., 2012; Miyoshi and Fishell, 2011; Rymar and Sadikot, 2007) and the mode of neurogenesis (Petros *et al*., 2015); all of which influence the cell cycle dynamics of VZ and SVZ cells. Additionally, many disease-associated genes are enriched in RGCs and interneuron progenitors (Schork et al., 2019; Trevino et al., 2020). Thus, a thorough characterization of gene expression in distinct neurogenic regions of the telencephalic VZ and SVZ will increase our understanding of initial cell fate decisions in the embryonic brain and provide insight into possible disease mechanisms.

In recent years, single-cell RNA-sequencing (scRNAseq) studies revealed extensive neuronal diversity in the mature telencephalon whereby cells can be classified into relatively clean distinct neuronal subtypes based on their transcriptome (Economo et al., 2018; Tasic et al., 2016; Tasic et al., 2018; Zeisel et al., 2015). However, significantly less is known about neural progenitor diversity within the developing embryo when initial fate decisions arise. Several initial studies performed bulk RNA-sequencing on the embryonic ventral forebrain, but these studies were not able to differentiate molecular characteristics between progenitor domains (Tucker et al., 2008; Zechel et al., 2014). More recent scRNAseq studies that targeted GEs found that initial signatures of mature interneuron (IN) subtypes appear in postmitotic precursors whereas very little transcriptional diversity was detected in VZ and SVZ progenitors (Chen et al., 2017; Mayer et al., 2018; Mi et al., 2018). However, these studies were underpowered for detecting potential transcriptional diversity in VZ neural progenitors because the GEs are comprised primarily of SVZ and mantle zone (MZ) cells at these mid-embryonic ages. Transcriptionally heterogeneous VZ cell populations exist in other central nervous system regions (Johnson et al., 2015; Li et al., 2020; Ogawa et al., 2005; Yuzwa et al., 2017). Thus, we still lack a definitive characterization of differential gene expression in VZ and SVZ neural progenitors in the embryonic mouse forebrain.

In this study, we performed scRNAseq analyses of LGE, MGE, CGE and CTX cells from E12.5 and E14.5 mice to identify spatial and temporal genetic heterogeneity of VZ and SVZ neural progenitors. To specifically increase the number of VZ cells, we utilized a reporter mouse line containing a destabilized VenusGFP protein that is driven by the *Nestin* promoter, which limits GFP leakiness into non-VZ cells and ensures GFP is a reliable readout of VZ cells (Sunabori et al., 2008). With this mouse line, we observed significant transcriptional diversity of VZ cells within the ventral telencephalon, both between different GEs as well as spatially segregated expression profiles of VZ cells within specific GEs. Many of these intriguing gene expression patterns were confirmed via multiplex *in situ* hybridizations. We also uncovered spatially-restricted gene expression patterns in the SVZ between and within the LGE, MGE, and CGE. Last, integrated transcriptional analysis of E12.5 and E14.5 highlighted differential gene expression between VZ cells during these embryonic timepoints, which could have important implications for the well-described temporal changes in interneuron cell fate (Bandler *et al*., 2017; Inan *et al*., 2012; Miyoshi and Fishell, 2011; Rymar and Sadikot, 2007). Thus, our data reveal both spatial and temporal transcriptional diversity in distinct VZ and SVZ progenitor cell populations which will further our understanding of neurogenesis during initial fate decisions in the embryonic forebrain.

## RESULTS

### Identification of distinct cell groups in the E12.5 mouse telencephalon

To characterize cellular heterogeneity in the developing telencephalon, we used the 10x Genomics scRNAseq platform to profile the transcriptome of cells from the MGE, LGE, CGE and CTX of E12.5 wild-type (WT) mice. At E12.5, neurogenesis has begun in the GEs, with most cells residing in the SVZ and MZ with significantly fewer cycling VZ neural progenitors (Turrero Garcia and Harwell, 2017; Wichterle et al., 2001). Conversely, cortical neurogenesis does not begin until ~E14 (Noctor et al., 2004; Sessa et al., 2008; Tyler et al., 2015), so the majority of E12.5 cortical cells are mainly VZ neural progenitors, specifically RGCs. To ensure that we collected a sufficient number of VZ cells from the GEs to identify transcriptional diversity in this population, we also dissected E12.5 brains from transgenic reporter mice that express destabilized Venus driven by the *Nestin* promoter and intronic enhancer (Nes-dVenus) (Sunabori *et al*., 2008) (Supplementary Figure 1A). We used fluorescence-activated cell sorting (FACS) to isolate GFP-positive cells from Nes-dVenus E12.5 mouse telencephalon (Supplementary Figure 1B). Note that ~87% of cortical cells are GFP-positive at E12.5 compared to 41-53% of GE cells, consistent with different proportions of VZ cells in these regions. After filtering out doublets and non-viable outliers (Supplementary Figure 2), we obtained >84,000 cells from the E12.5 mouse brain (~24,000 from WT mice and ~60,000 from the Nes-dVenus mice) that were used for subsequent analysis (Figure 1A).

**Figure 1.**
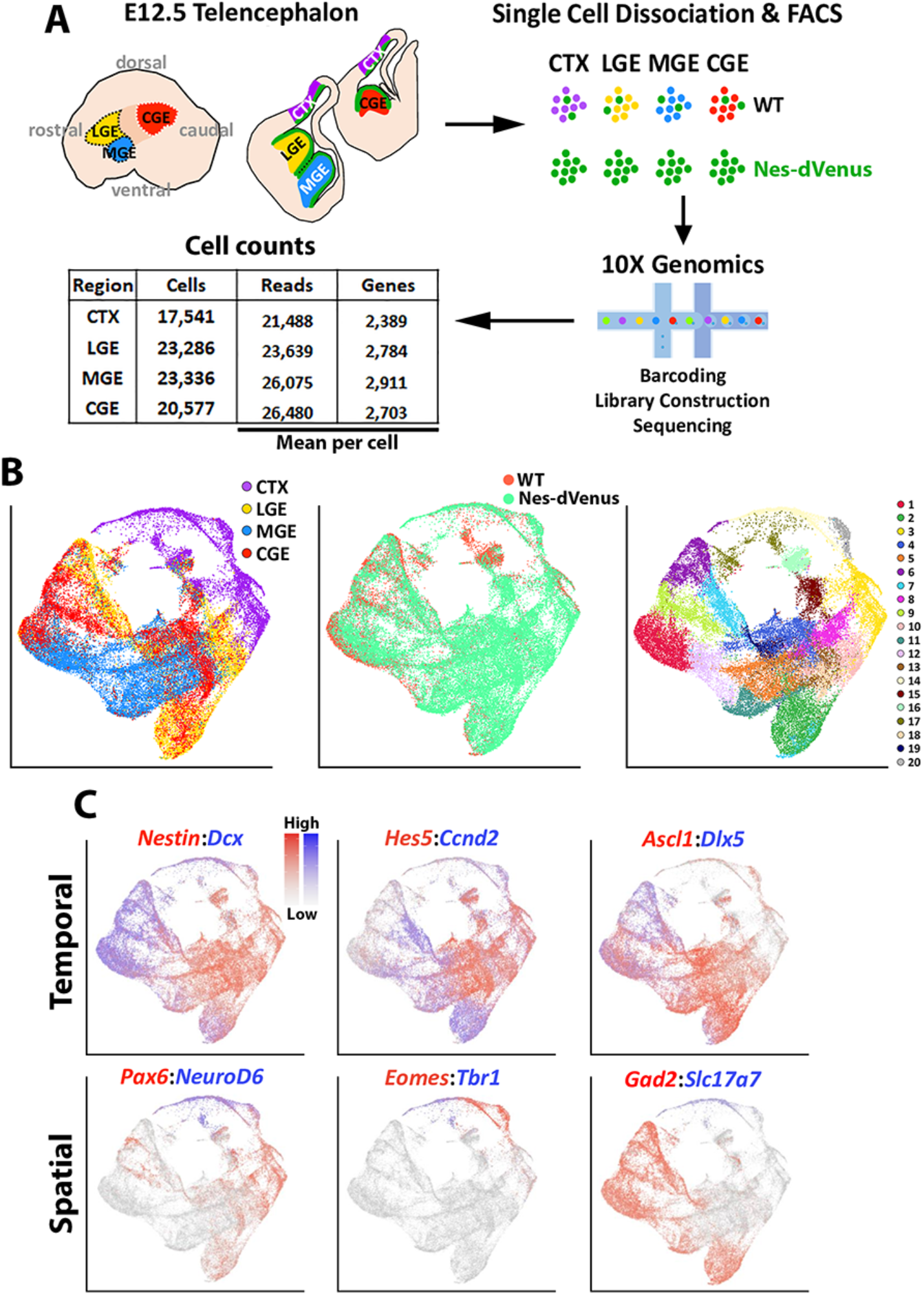
scRNAseq of distinct E12.5 forebrain regions from WT and Nes-dVenus mice. (A) Schematic of telencephalic cell dissection, isolation and 10X Genomics workflow. Chart depicts the total number of combined WT and Nes-dVenus sequenced cells from each embryonic brain regions, including the mean reads per cell and genes per cell. (B) Uniform manifold approximation and projection (UMAP) plots of all cells labeled by brain region (left), mouse line (middle) or putative cell cluster annotation (right). (C) Representative genes displaying enriched temporal and/or spatial expression patterns. CTX, cortex; LGE, lateral ganglionic eminence; MGE, medial ganglionic eminence; CGE, caudal ganglionic eminence; WT, wild-type. See also Supplementary Figures 1-3.

We used the Seurat software package (Stuart et al., 2019) to integrate these cells and identified 20 cell clusters that were largely segregated by brain regions, with clean separation of the CTX and MGE cells while there was greater overlap between the LGE and CGE populations (Figure 1B). We observed common VZ markers (*Nestin* and *Hes5*) and SVZ/early neurogenic markers (*Dcx* and *Ccnd2*) present in cells from all 4 regions, as well as GABAergic-specific (*Ascl1, Dlx5*, and *Gad2*) and glutamatergic-specific (*Pax6, Neurod6, Eomes, Tbr1*, and *Slc17a7*) genes restricted to their expected populations (Figure 1C). By extracting each region individually from the dataset, we observed many genes strongly enriched in the MGE, LGE, CGE and CTX. While some of these genes have well-described roles in specific brain regions, we did uncover many genes whose regional restricted patterns have not been previously described, such as the alpha-internexin encoding gene *Ina* and Insulin like Growth Factor Binding Protein 5 encoding gene *Igfbp5* in the CGE (Supplementary Figure 3). Together, these data reveal different cohorts of telencephalic cells display distinct region-specific gene expression profiles exist at E12.5.

### Common neurogenic zones within each ganglionic eminence

To identify specific cell clusters in the GEs, we removed cortex-derived cells and identified eighteen GE-derived cell clusters (Figure 2A). Based on *Nestin* and *Dcx* expression patterns, we find eight clusters that likely represent VZ cells (high *Nestin*), six clusters that likely represent postmitotic cells (high *Dcx*), and four clusters that likely represent SVZ basal progenitors (moderate levels of both *Nestin* and *Dcx*). To confirm that we successfully enriched for *Nestin*-expressing VZ cells with the Nes-dVenus mouse, we plotted GE cells based on which mouse line they were derived from. This approach clearly demonstrates that the vast majority of high *Nestin*-expressing cells are indeed derived from the Nes-dVenus mouse, with very few VZ cells captured in the WT mice (Figure 2A). Violin plots revealed that most cells expressing VZ-enriched genes such as *Nestin, Hes5* and *Ccnd1* were harvested from the Nes-dVenus mouse whereas a greater percentage of cells expressing more mature markers such as *Dcx, Tubb3* and *Gad2* arose from the WT mice (Supplementary Figure 4A). Thus, we have a significantly greater percentage of VZ cells compared to previous studies (Mayer *et al*., 2018; Mi *et al*., 2018) (Supplementary Figure 5), which provides us greater power to identify transcriptional heterogeneity and smaller subgroups in this VZ cell population that may have been previously overlooked.

**Figure 2.**
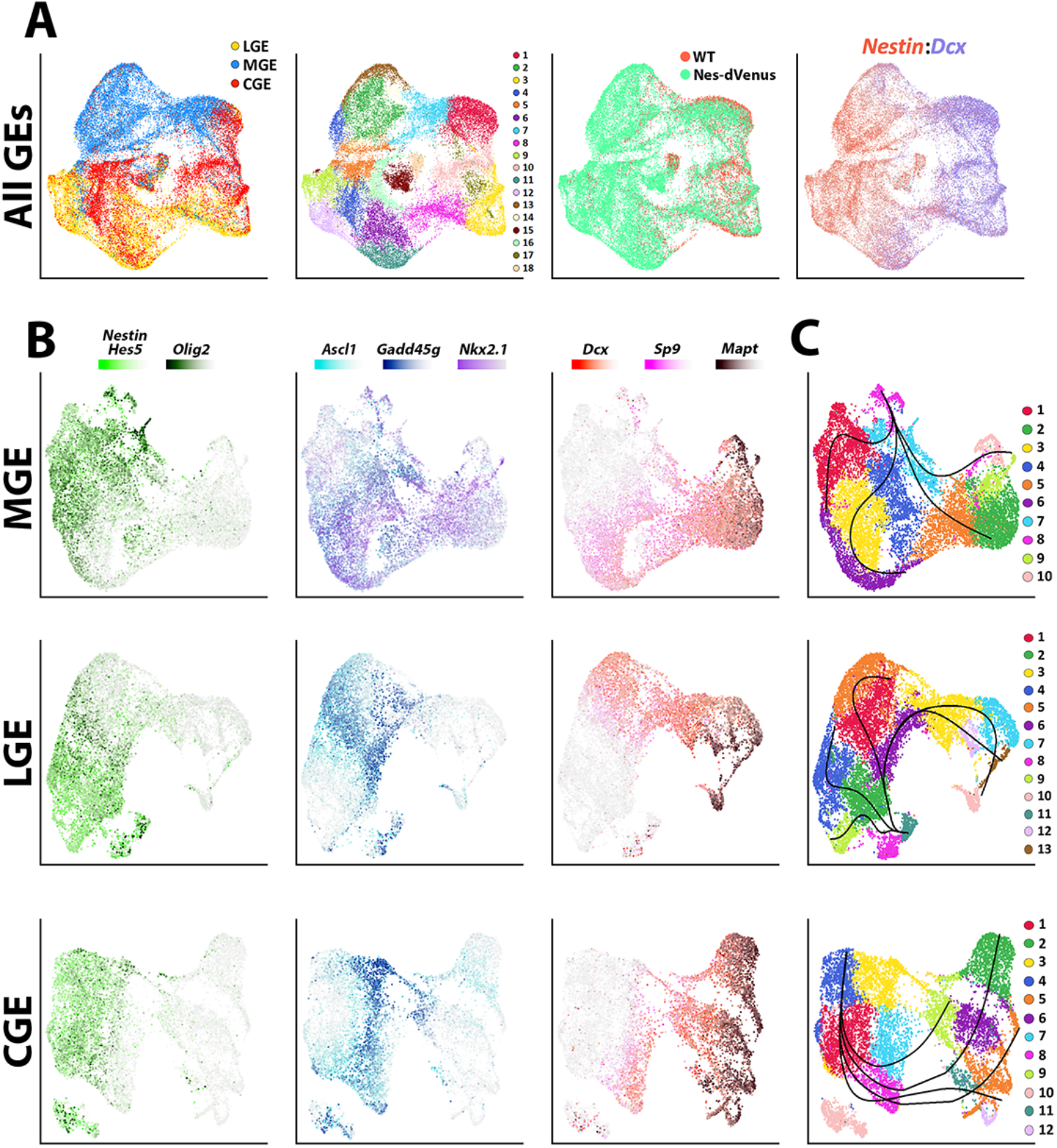
Common neurogenic lineages in each ganglionic eminence. (A) UMAP plots displaying all GE-derived cells annotated from left to right by: brain region, putative cell cluster annotation, mouse strain, and expression of VZ marker *Nestin* and SVZ/MZ marker *Dcx*. (B) UMAP visualizations show transcriptional diversity in each GE region. Several genes are depicted to highlight distinct neurogenic stages: *Nestin, Hes5* and *Olig2* are indicative of VZ cells (green), *Asc1* and *Gadd45g* labeling intermediate SVZ cells (blue), and *Dcx, Sp9* and *Mapt* depicting postmitotic neuronal precursors (red). (C) UMAP plots of MGE, LGE and CGE cells annotated via putative cell clusters and including Slingshot analyses depicting developmental progression through neurogenic stages. See also Supplementary Figures 4-6.

The LGE, MGE and CGE cells were individually extracted out and reclustered to reveal three common cell types shared between regions: 1) *Nestin, Hes5*, and *Olig2* expressing mitotic VZ cells, 2) *Ascl1, Ccnd2*, and *Gadd45g* expressing SVZ/BP cells, and 3) *Dcx, Sp9*, and *Mapt* expressing postmitotic SVZ and MZ cells (Figure 2B and Supplementary Figure 6). Focusing on MGE-specific genes, *Nkx2.1* was expressed by most cells within the MGE whereas *Lhx6* was restricted to postmitotic MGE cells, a subset of which also expressed the mature MGE-derived subtype marker *Sst* (Supplementary Figures 4B and 6). Within LGE and CGE clusters, *Ebf1*-positive, *Mapt*-positive and *Sp9*-negative clusters likely identify GABAergic projection neurons (Nery et al., 2002) (Supplementary Figure 6). Also, *Npy*- and *Sst*-positive clusters were captured within LGE and CGE dataset representing MGE-derived migrating cortical interneurons migrating through both regions towards the cortex (Kessaris et al., 2014) (Supplementary Figure 6). Pseudotime analysis with Slingshot (Street et al., 2018; Trapnell et al., 2014) showed clear trajectories in each region originating from *Nestin*- and *Hes5*-positive clusters progressing towards the postmitotic cell markers (Figure 2C). This analysis displays many branching patterns within VZ neural progenitors which could reflect distinct developmental trajectories among *Nestin*-positive VZ cells in GEs.

We next asked how GE-derived *Nestin*-expressing VZ neural progenitors are transcriptionally different from cycling *Dcx*-expressing SVZ cells and postmitotic SVZ/MZ cells.

To specifically extract the VZ cells, we isolated cells that had a *Nestin* expression value above 1.5 from the log normalized GE count data (high *Nestin* cells). The value of this threshold was optimized by testing different expression values for subsetting, with the goal of harvesting a large number of high *Nestin*-expressing cells with minimal contamination of *Dcx*-expressing cells (Supplementary Figures 7A-B). To obtain a relatively clean population of cycling SVZ/BP cells, we isolated cells that expressed *Dcx, Mki67* and *Ccnd2* but were negative for *Mapt* and *Rbfox3*. To enrich for postmitotic SVZ/MZ cells, we isolated cells that were negative for *Nestin, Hes5, Mki67, Ccnd1*, and *Ccnd2* but were positive for *Dcx* (Figure 3A). The subsetting value for postmitotic cells (high *Dcx* cells) was optimized with the similar approach to that of *Nestin*-expressing cell enrichment (Supplementary Figures 7C-D). These three distinct cell populations retained their regional segregation and were grouped into 12 clusters (Figure 3A). Comparative analysis between the *Nestin*-positive, mitotic *Dcx*-positive, and postmitotic *Dcx*-positive groups showed clear transcriptional diversity, with 430 genes enriched in VZ cells, 187 genes upregulated in mitotic SVZ cells, and 370 genes upregulated in postmitotic SVZ/MZ cells (minimum log-fold change = 0.25) (Table 1). Highlighting the top 20 enriched genes for each cell type revealed several neural stem cell-associated genes for VZ cells (*Fabp7, Hes5, Mt3, Ccnd1*, and *Ptprz1*) and BP-associated genes for cycling SVZ cells (*Abracl, Mpped2, Dlx6os1, Ccnd2*, and *Sp9*) whereas mature neuronal makers (*Mapt, Ina, Tubb2a, Rtn1*, and *Tubb3*) were detected in the postmitotic SVZ/MZ cells (Figure 3B-C). Each of the 12 identified cell clusters contained several enriched genes that correlated with their cell types, demonstrating GE-specific gene expression within each VZ, SVZ and MZ domain (Figure 3D). Our data uncovers transcriptional diversity within distinct neurogenic domains that may provide insight into how specific genes regulate early cell fate decisions within the embryo.

**Figure 3.**
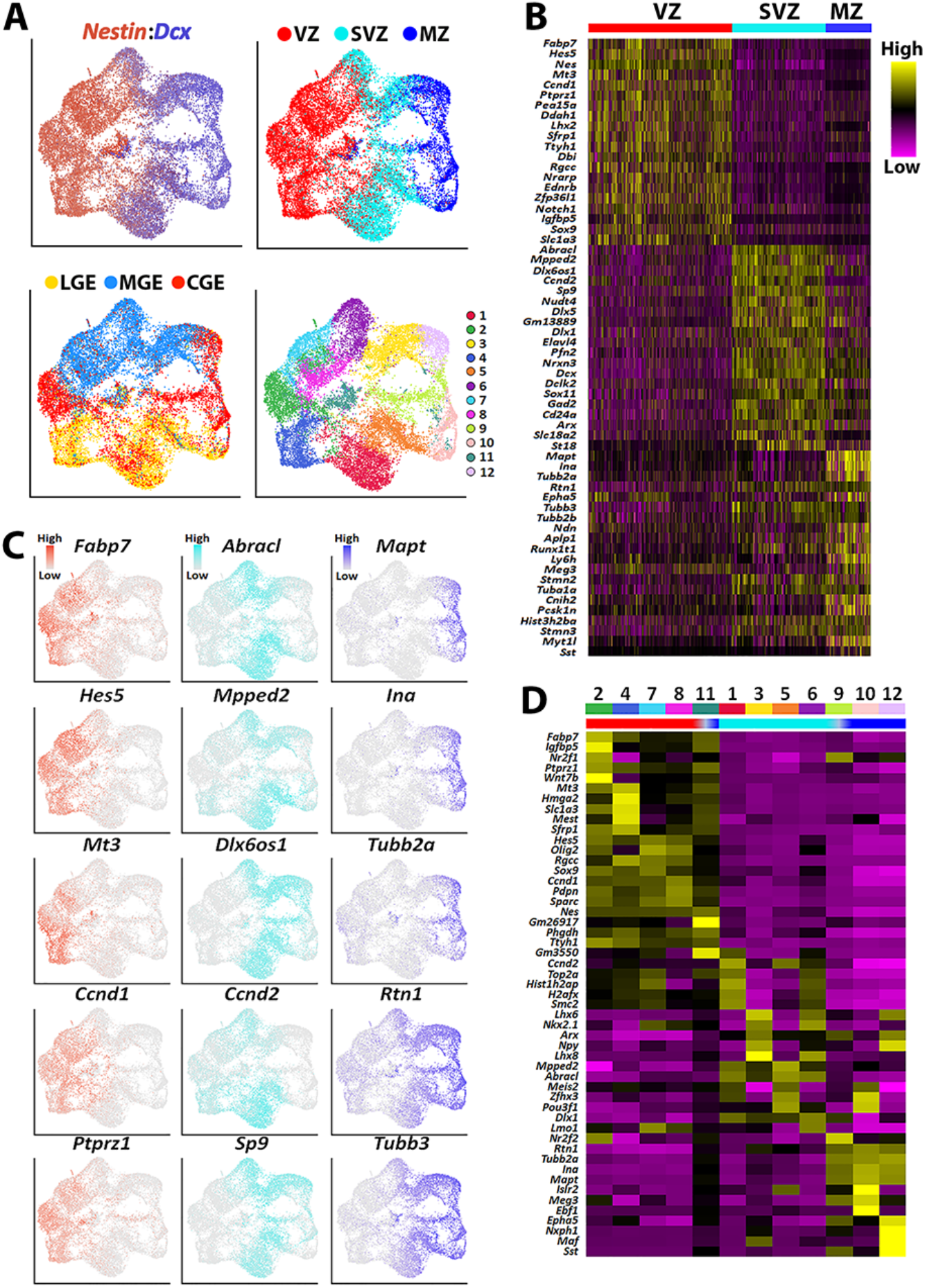
Transcriptional heterogeneity within the VZ, SVZ and MZ in E12.5 GEs. (A) High *Nestin*- and *Dcx*-expressing cells (>1.5 fold-expression above normalized dataset) were extracted from the GE populations and replotted. UMAP plots are annotated by: expression of *Nestin* and *Dcx* (upper left), brain region (lower left), separated into VZ (high-*Nestin*, red), SVZ (*Dcx-, Mki67*-, and *Ccnd2*-expressing, light blue) and MZ (*Dcx*-positive and *Ccnd2*- and *Mki67*-negative, dark blue) (upper right), and putative cell clusters (lower right). (B) Heatmap of genes enriched in VZ, SVZ and MZ cells. Each column represents expression in a single cell, color-coded as per the color scale. (C) UMAP plots depicting the top 5 differentially expressed genes (DEGs) in the VZ, SVZ and MZ regions. (D) Heatmap showing expression of top 5 DEGs in each cell cluster from (A), with colored bar depicting whether each cluster contains predominantly VZ (red), SVZ (light blue) or MZ (dark blue) cells.

### Transcriptional heterogeneity throughout the ventricular zone

To uncover transcriptional heterogeneity specifically in VZ cells, we extracted and replotted the high *Nestin*-expressing cells (threshold ≥ 1.5) from the LGE, MGE and CGE (Figure 4). Cells were largely clustered based on brain region and were grouped into 10 clusters. There was a large cohort of VZ-enriched genes that were restricted to one specific GE (Figure 4B-C), with some genes being confined to specific VZ clusters (subdomains) within a GE (Figure 4D). We selected ~20 genes that displayed intriguing spatial or cell type-specific expression patterns within the VZ, the vast majority of which have not been previously described, and visualized them with the RNAscope HiPlex *in situ* hybridization assay. We identified several pan-VZ genes such as *Ednrb, Pkdcc* and *Nrarp* that are known to regulated by Wnt and Notch signaling pathways that are critical for embryonic development (Krebs et al., 2001; Takeo et al., 2016; Vitorino et al., 2015), as well as mitosis associated genes *Prc1, Cenpf*, and *Ube2c* (Engeland, 2018) (Figure 5A-C and Supplementary Figure 7). There were also genes that were strongly expressed in VZ cells in only 2 regions: *Ptx3* was expressed in VZ cells throughout the LGE and CGE yet absent from the MGE, while *Fgfr3* was strongly expressed in MGE and CGE VZ cells and absent in the LGE (Figure 5D; Supplementary Figures 4C, 6C and 7). And several genes were restricted to VZ cells in only one region, such as *Shisa2* and *Cntnap2* in the LGE, *Igfbp5* in the CGE, and *Asb4* in the MGE (Figure 5E and Supplementary Figures 4C, 6C and 7).

**Figure 4.**
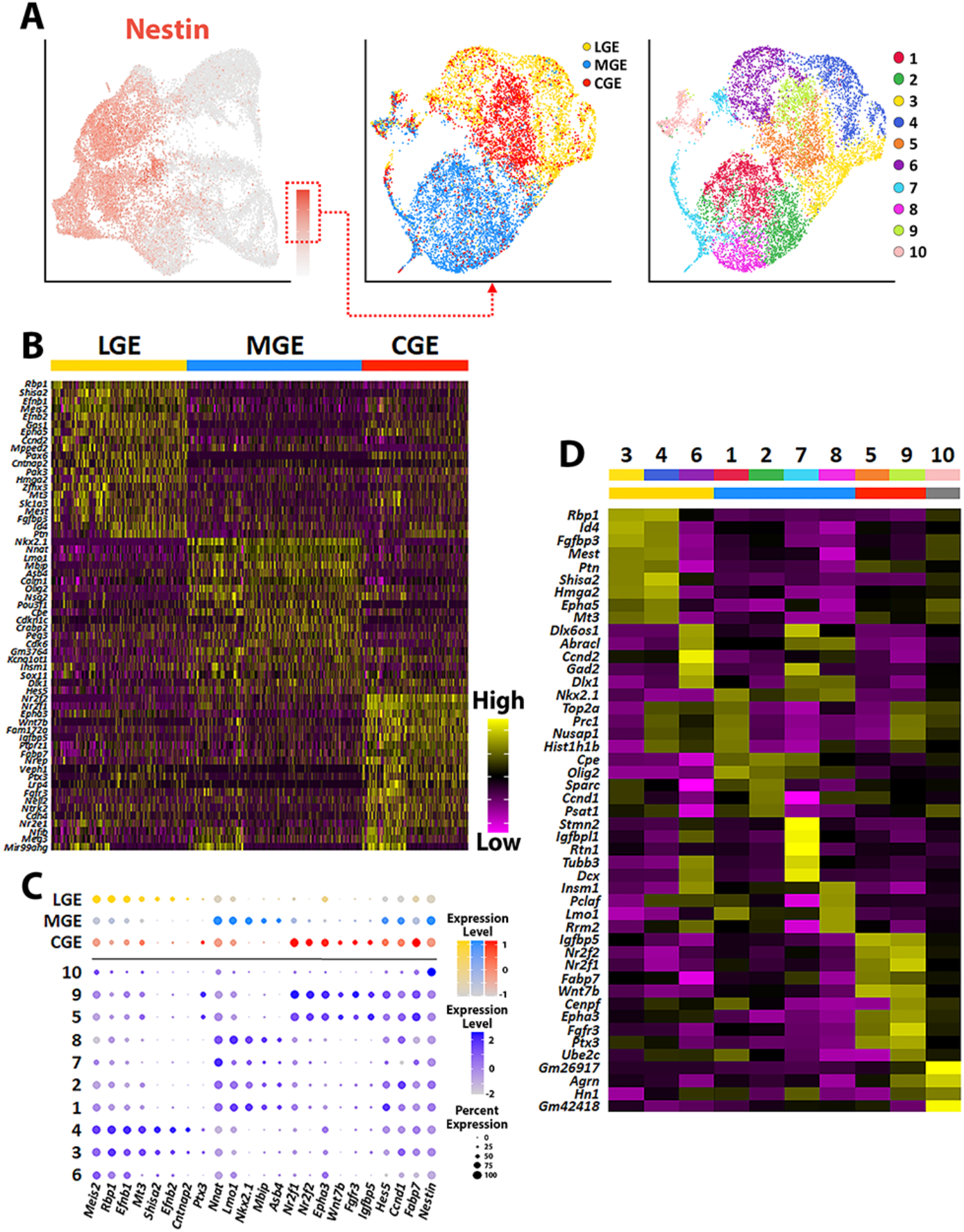
Transcriptional heterogeneity of *Nestin*-expressing VZ cells in the ganglionic eminences. (A) High *Nestin*-expressing (>1.5 normalized dataset) cells were extracted from GE dataset and replotted as UMAP graphs annotated by brain region (middle) and putative cell clusters (right). (B) Heatmap of top 20 DEGs enriched in VZ cells from the MGE, LGE and CGE. Each column represents expression in a single cell, color-coded as per the color scale. (C) Dot-plot depicting expression levels of DEGs within each brain region (MGE, LGE and CGE) and cell cluster. (D) Heatmap showing expression of top 5 DEGs in each cluster from (A), with colored bar depicting whether each cluster contains cells from the LGE (yellow), MGE (blue) or CGE (red). Gray bar indicates cluster containing cells from all three GEs. Each column represents averaged expression in cells, color-coded as per the color scale. See also Supplementary Figure 7.

**Figure 5.**
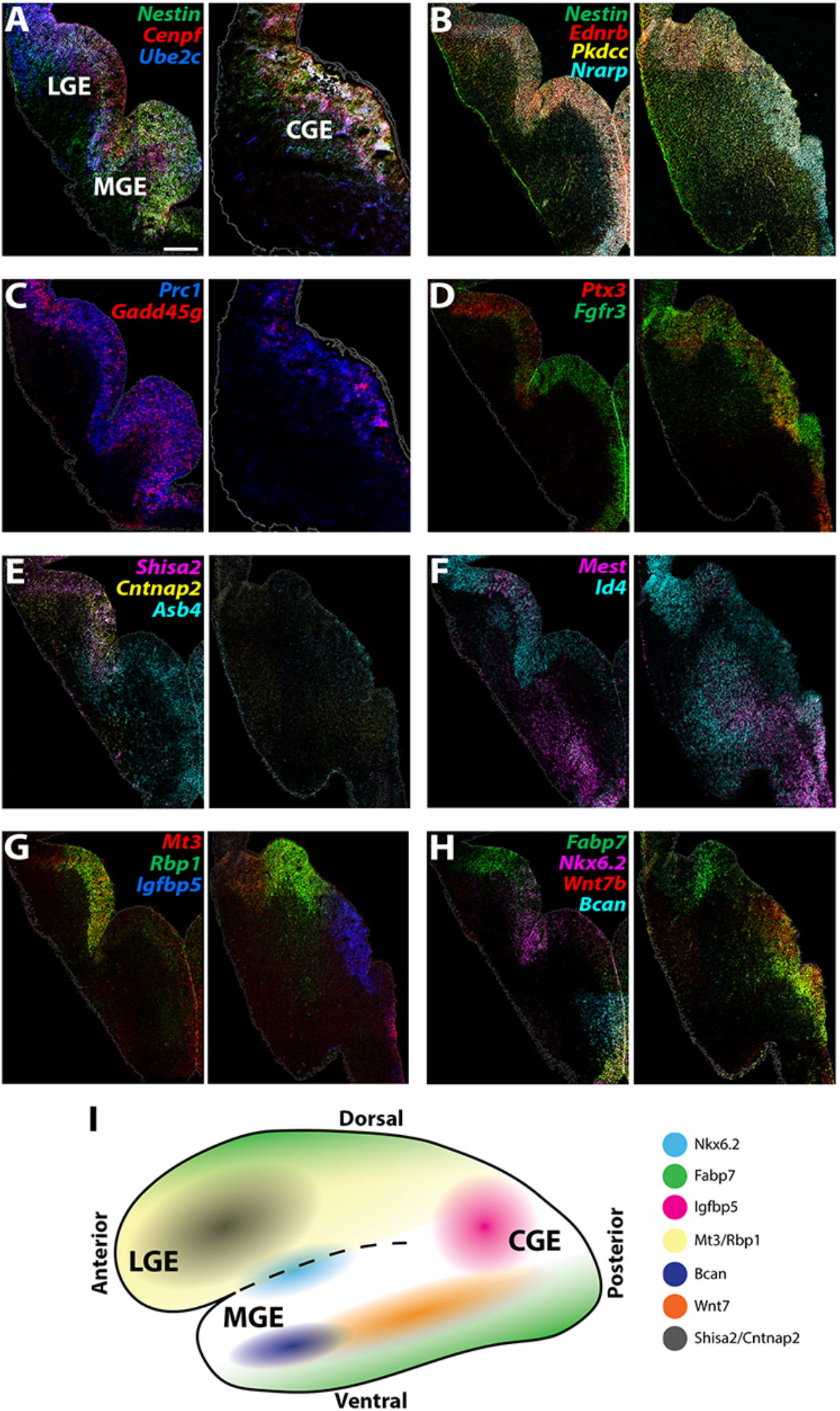
Expression profiles of spatially-enriched genes in VZ cells in the ganglionic eminences. (A-H) RNAscope HiPlex and HiPlexUp assays of differentially expressed VZ genes on E12.5 GE coronal sections. Pan-VZ genes are shown in panels A-B while more restricted expression patterns are displayed in panels C-H. Scale bar in A = 200 μm. (I) Schematic of a whole mount view of the E12.5 ventral telencephalon depicting the approximate spatial location of VZ-enriched genes within the GEs. See also Supplementary Figure 8.

We also observed many genes that had more refined spatially restricted expression patterns within various GEs, highlighting several examples below. *Gadd45g*, a gene that regulates stem cell proliferation (Kaufmann and Niehrs, 2011), displayed a salt-and-pepper profile with enriched expression in the dorsal LGE (dLGE) and ventral MGE and CGE (vMGE and vCGE) (Figure 5C; Supplementary Figure 7). *Id4* and *Mest* displayed similar expression profiles in their dot plots (Supplementary figure 7). However, *Id4* was detected in VZ cells throughout the LGE, CGE and dMGE, whereas *Mest* was restricted specifically to VZ cells in the dLGE (Figure 5F; Supplementary Figure 7). *Id4* and *Mest* were both enriched in the SVZ/MZ of the vMGE and CGE, with a bias for *Id4* to the dCGE and *Mest* to the vCGE, but only *Id4* was present in the SVZ layer of the LGE. These distinct differences highlight the need to confirm transcriptome expression profiles from scRNAseq experiments via *in situ* hybridizations.

*Mt3* and *Rbp1* are both strongly enriched in middle-to-ventral LGE VZ cells and in the dCGE, with only a smattering of *Mt3*- and *Rbp1*-expressing cells observed in the MGE (Figure 5G; Supplementary Figures 4C, 6C and 7). *Igfbp5* expression is exclusively restricted to the vCGE VZ cells (Figure 5G; Supplementary Figures 4C, 6C and 7). *Bcan* and *Wnt7b* are enriched in vMGE VZ cells with *Wnt7b* expression extends into the vCGE, whereas *Nkx6.2* is expressed at the LGE/MGE boundary as previously described (Sousa et al., 2009) (Figure 5H; Supplementary Figures 4C, 6C and 7). Of note, *Fabp7* is strongly enriched in VZ cells in the dLGE, vMGE, and both dCGE and vCGE, as if forming longitudinal stripes along the dorsal and ventral GE boundaries (Figure 5H and Supplementary Figures 4C and 7). In sum, our data reveal previously unidentified transcriptional heterogeneity within VZ cells throughout the GEs (Figure 5I), with many gene expression patterns characterized here for the first time.

### Transcriptional heterogeneity in SVZ/MZ cells within the ventral telencephalon

We utilized a similar approach to extract SVZ/MZ cells from the GEs that had high expression levels of *Dcx*, which displayed fairly clean segregation between LGE, MGE and CGE resulting in 11 cell clusters (Supplementary Figure 8A). We observed clusters containing previously described region-enriched genes such as *Meis2* in the LGE, *Nkx2.1, Lhx6* and *Lhx8* in the MGE, and *Nr2f1* and *Nr2f2* for CGE (Kanatani et al., 2008; Lodato et al., 2011) (Supplementary Figures 8B-C and 9); the expression profile of these genes was confirmed with *in situ* hybridizations (Figure 6A-B, G). Of note, *Nr2f1* and *Nr2f2* are also enriched in the dLGE and POA, in agreement with previous observations (Figure 6B and Supplementary Figure 4B) (Hu et al., 2017). We also identified markers that separated cycling SVZ/BP cells from postmitotic MZ cells. Genes such as *Ascl1, Dlk2, E2f1* and *Tcf4* are expressed in the VZ-SVZ boundary throughout all GEs and then downregulated in postmitotic MZ cells (Figure 6C-D; Supplement Figure 9). Conversely, *Zfhx3, Ina, Mapt, Mpped2* and *Stmn2* are predominantly expressed in *Dcx*-positive postmitotic cells in the MZ layers, with *Zfhx3* and *Mpped2* expressed near the SVZ-MZ boundary whereas *Ina, Mapt* and *Stmn2* are found in deeper MZ regions (Figure 6D-G; Supplement Figure 9).

**Figure 6.**
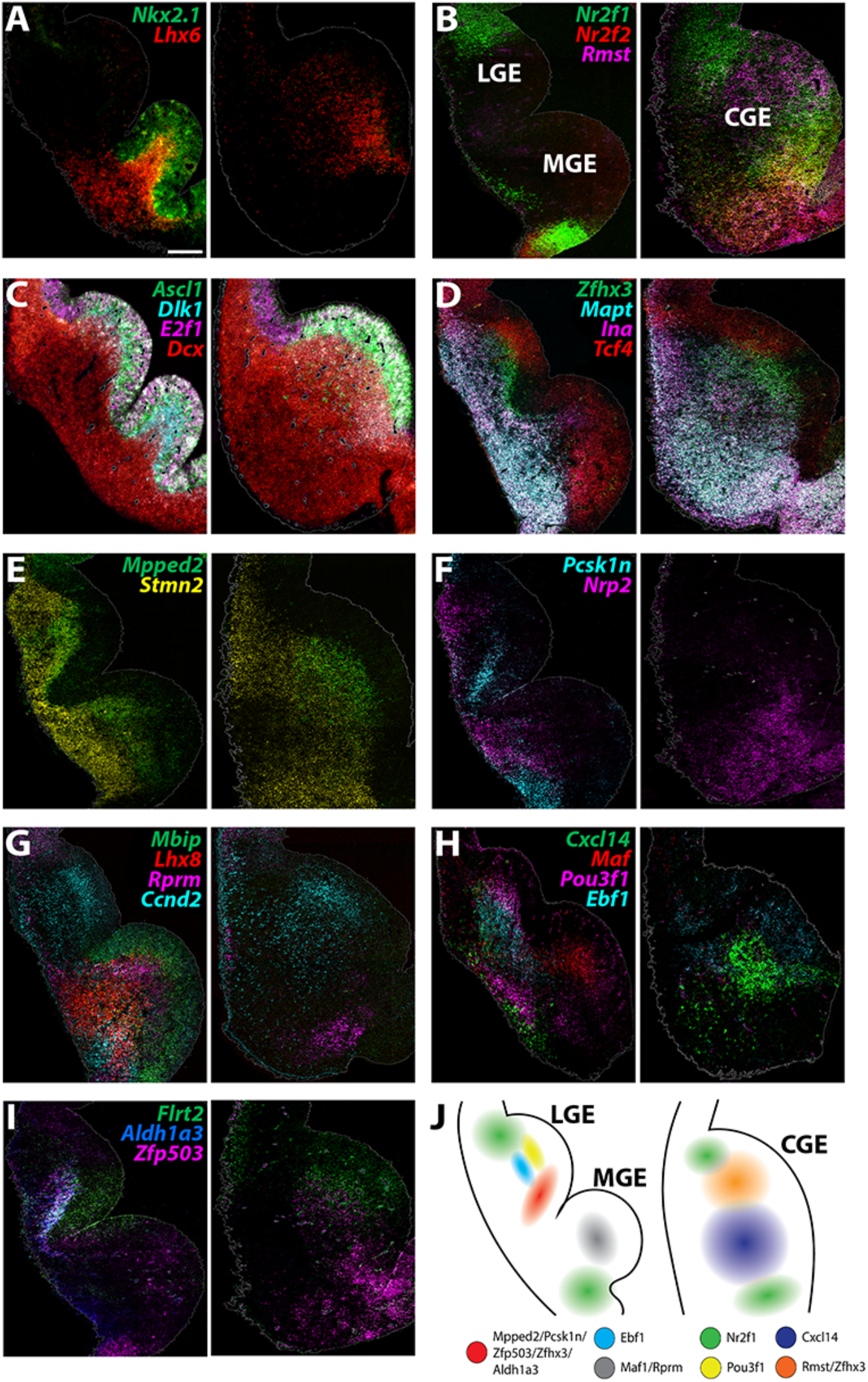
Expression profiles of spatially-enriched genes in SVZ/MZ cells in the ganglionic eminences. (A-I) RNAscope HiPlex and HiPlexUp assays of differentially expressed SVZ/MZ genes on E12.5 GE coronal sections. Scale bar in A = 200 μm. (J) Schematic of a coronal section through the E12.5 ventral telencephalon depicting the approximate spatial location of SVZ/MZ-enriched genes within the GEs. See also Supplementary Figures 9-10.

Other markers displayed more restricted expression patterns within the GEs. For example, *Nrp2* is enriched in deep MZ cells within the dLGE, dMGE and vCGE whereas *Pcsk1n* displays a complementary pattern in the vLGE and vMGE but is not expressed within the CGE (Figure 6F). *Ccnd2* and *Flrt2* were enriched in the SVZ layer of both LGE and CGE with significantly weaker expression in the MGE (Figure 6G & I and Supplement Figure 9). *Aldh1a3* and *Zfp503* are strongly enriched in MZ cells in the vLGE, with *Zfp503* expression extending into the SVZ/MZ layers of the vCGE (Figure 6I; Supplement Figure 9). *Mbip* and *Rprm* are both enriched in the MGE SVZ, with *Mbip* located along the VZ/SVZ border and *Rprm* at the SVZ/MZ border (and *Rprm* is also found in the vCGE) (Figure 6G). *Rmst* and *Cxcl14* were restricted to the CGE, with *Rmst* enriched in the dCGE (somewhat complementary to *Nr2f1/2*) and *Cxcl14* found in deeper MZ cells (Figure 6B & G). In a similar spatial relationship, *Pou3f1* and *Ebf1* are restricted to the LGE, with *Ebf1*-expressing cells located slightly deeper to *Pou3f1*-expressing cells at the SVZ/MZ border (Figure 6H). These single cell gene expression profiles provide new insight into the transcriptional diversity in *Dcx*-positive SVZ/MZ postmitotic GE cells (Figure 6J) and revealed subdomains of genetically defined SVZ/MZ cells that have not been previously described.

### Developmental Transition of Ventral Telencephalic VZ Neural Progenitors Over Time

There is ample evidence that the capacity for neural progenitors to generate specific neuronal subtypes changes over time (Gal et al., 2006; Pilz et al., 2013). In the MGE, production of SST-expressing interneurons is significantly decreased at E14.5 compared to E12.5 (Bandler *et al*., 2017; Inan *et al*., 2012; Miyoshi and Fishell, 2011). To characterize transcriptional changes in GE progenitors over time, we collected LGE, MGE and CGE cells from E14.5 WT and Nes-dVenus mice. These cells were largely segregated by brain region and consisted of 19 cell clusters (Supplement Figure 10A-C). As with the E12.5 population, significant heterogeneity was still observed when extracting out the highest *Nestin*- and *Dcx*-expressing cells (Supplementary Figure 10D-G).

To compare the E12.5 and E14.5 cells, we integrated these datasets together. The majority of postmitotic cell clusters consisted of E14.5 cells, and most *Nestin*-expressing cells were derived from the Nes-dVenus mouse (Figure 7A). These integrated populations were still primarily segregated based on region and were divided into sixteen different clusters (Figure 7B-C). We isolated strong *Nestin*- and *Dcx*-expressing cells from the integrated dataset and performed a differential expression analysis on both sets of cells (Figure 7D-F). High *Dcx*-expressing from E12.5 and E14.5 were nearly completely overlapping with no significant differences, indicating that the global population of postmitotic cells from GEs are transcriptionally very similar at these ages (Figure 7F). However, when we compared the high *Nestin*-expressing cells, we observed more age-specific segregation in the dot plot clusters, with one subdomain predominately consisting of E12.5 cells and another population containing E14.5 cells (Figure 7E).

**Figure 7.**
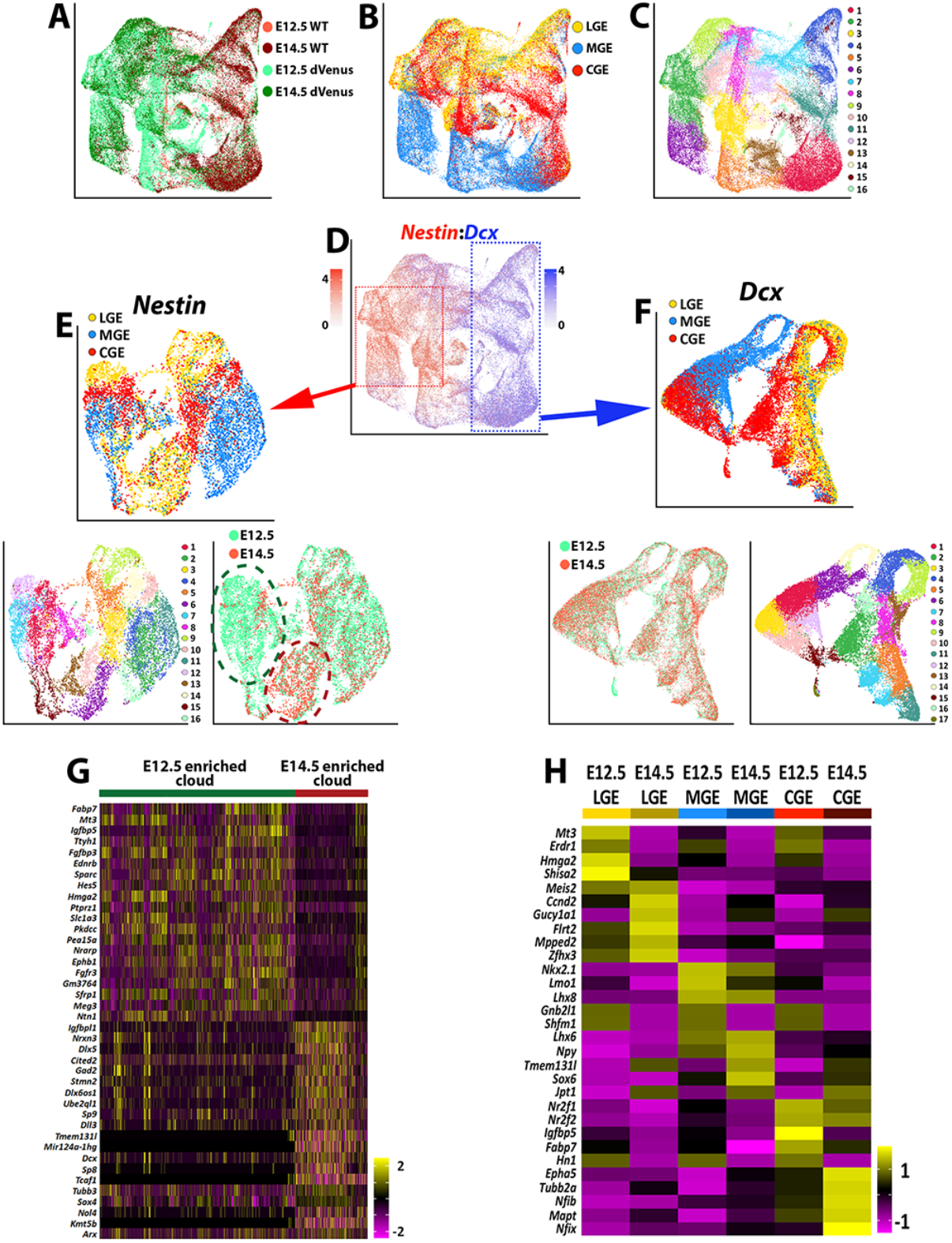
Comparison of transcriptional changes between E12.5 and E14.5 ventral telencephalon. (A-C) UMAP plots of all E12.5 and E14.5 cells annotated by mouse line and embryonic timepoint (A), brain region (B), and putative cell cluster (C). (D) UMAP plot of E12.5 and E14.5 cells depicting expression levels of *Nestin* and *Dcx*. Red box = approximate location of high *Nestin*-expressing cells (>1.5 normalized dataset), blue box = approximate location of high *Dcx*-expressing cells (>1.5 normalized dataset). (E) UMAP plot depicting high *Nestin*-expressing cells annotated by brain region (top), putative cell clusters (lower left) and timepoint (lower right). Note the clouds that are strongly enriched for E12.5 cells (dark green oval) and E14.5 cells (dark red oval). (F) UMAP plot depicting high *Dcx*-expressing cells annotated by brain region (top), putative cell clusters (lower right) and timepoint (lower left). Note that E12.5 and E14.5 cells are largely overlapping populations with no clear differential clustering. (G) Heatmap depicting the top 20 DEGs between the E12.5-enriched (dark green oval) and E14.5-enriched (dark red oval) clouds of VZ cells from (E). The E14.5-derived cells were removed from the E12.5-enriched cloud, and vice versa. Each column represents expression in a single cell, color-coded as per the color scale. (H) Heatmap depicting the top 5 DEGs in VZ cells from each GE region at E12.5 and E14.5. Each column represents averaged expression in cells, color-coded as per the color scale. See also Supplementary Figures 11-12.

To compare these two cell clusters, we removed the E12.5 cells from the E14.5 cloud (and vice versa) and identified many genes enriched specifically in E12.5 or E14.5 *Nestin*-expressing cells (Figure 7G). Changes in VZ gene expression between these two timepoints were even observed within specific GEs (Figure 7H). Many genes enriched in E14.5 high *Nestin*-expressing are commonly associated with BPs or postmitotic cells such as *Dcx, Tubb3, Mpped2, Arx* and *Nrxn3* (Figure 7G). The adult neural progenitor marker *Igfbpl1* (Artegiani et al., 2017) was also upregulated in E14.5 VZ cells when compared to E12.5 (Figure 7G). To ensure that the dVenus is not ‘leaking’ into non-VZ cells at E14.5 and contaminating this dataset with SVZ/BP cells, we confirmed that many previously reported pan-VZ cell markers are still expressed in the E14.5 high *Nestin*-expressing cells (Supplementary Figure 11). Taken together, this data suggests that while *Dcx*-positive cells are transcriptionally very similar between E12.5 and E14.5, *Nestin*-positive VZ cells in the GEs display a transition in their gene expression profiles, with a greater number of neurogenic and postmitotic genes present in E14.5 VZ cells.

## DISCUSSION

We performed scRNAseq on distinct neurogenic forebrain regions to generate a comprehensive transcriptional overview of embryonic telencephalic progenitors. By utilizing the Nes-dVenus mouse, we were able significantly increase the number of VZ cells and differentiate between cycling VZ cells (RGCs/APs) and cycling SVZ cells (BPs), a distinction that had not been explored in previous forebrain scRNAseq studies. This approach revealed a rich diversity of gene expression profiles in progenitors throughout the VZ and SVZ/MZ of the ventral forebrain. These insights could increase our understanding of GABAergic neuronal cell types patterning during neurogenesis and have important implications for understanding neurodevelopmental disorders because many disease-associated genes are enriched in radial glia/VZ cells and interneuron progenitors (Schork *et al*., 2019; Trevino *et al*., 2020).

We uncovered striking spatial transcriptional diversity throughout the VZ layers, with many genes displaying spatially-restricted expression patterns that had not been previously reported. Some genes were restricted to one or two GEs, with several genes restricted or strongly enriched in the CGE. There are minimal tools to genetically target CGE-specific cell types during embryogenesis because of the lack of established CGE-specific genes, and even some traditionally described CGE-specific genes such as *Nr2f1* and *Nr2f2* are also expressed in the posterior MGE (Hu *et al*., 2017). Our finding that *Igfbp5* is restricted to the CGE at E12.5 (Figure 5 and Supplementary Figure 7) warrants further investigation into *Igfbp5* as a promising tool to label CGE-derived cells.

Numerous genes display restricted expression patterns that were enriched in dorsal or ventral VZ cells in specific structures. This is particularly relevant due to the ample evidence that certain spatial domains are biased to generate particular interneuron subtypes (Brandao and Romcy-Pereira, 2015; Flames *et al*., 2007; McKinsey *et al*., 2013; Tyson *et al*., 2015; Wonders *et al*., 2008; Xu *et al*., 2010). While a relationship between spatial domain and interneuron subtype has been described in the MGE, whether distinct CGE-derived interneuron subtypes arise from specific CGE regions remains unknown. Several genes such as *Rbp1, Id4* and *Wnt7b* are enriched in subdomains of VZ cells within the CGE, which may indicate that certain CGE domains preferentially give rise to specific interneuron subtypes. Another intriguing expression profile was *Fabp7*, which is traditionally believed to be a pan-VZ/radial glia cell marker (Arai et al., 2005; Schmid et al., 2006; Yun et al., 2012; Yuzwa *et al*., 2017). However, in the ventral telencephalon, *Fabp7* is restricted to the dorsal and ventral GE boundaries, as if demarcating the GEs from the surrounding forebrain.

We also detected transcriptionally defined subdomains within the SVZ/MZ layers to a greater extent than was previously appreciated. For example, *Mbip* and *Rprm* show a complimentary expression profile at the VZ/SVZ and SVZ/MZ border in MGE, respectively. We observed several genes (*Mpped2, Pcsk1n, Aldh1a3, Zfhx3* and *Zfp503*) strongly enriched at the SVZ/MZ boundary in the ventral LGE (Figure 6). Within the CGE, *Nrp2, Rprm* and *Zfp503* are enriched in the ventral region whereas *Mpped2, Ccnd2* and *Flrt2* are enriched in the SVZ/MZ of the dorsal CGE. As cell fates become more defined as cells transition from cycling to postmitotic at this SVZ/MZ border, further work is needed to characterize the role of these spatially restricted genes in this process.

We also identify a transition in the transcriptional profile of VZ cells from E12.5 to E14.5 that was not observed in SVZ/MZ cells, which were transcriptionally very similar between the two ages. In general, E14.5 VZ cells from the GEs begin to express genes that are indicative of more mature cells, such as *Dcx, Arx* and *Gad2*. These transcriptional changes are occurring during a simultaneous increase in the number of SVZ cells/BPs with a proportional decrease in the number of proliferative VZ cells, as most MGE VZ cells give rise to SVZ/BP cells by E14.5 (Glickstein et al., 2007; Petros *et al*., 2015; Tsoa et al., 2014). Gene expression changes in VZ cells over time could guide this transition in cell cycle dynamics and in part regulate temporal changes in cell fate throughout neurogenesis (Bandler *et al*., 2017; Inan *et al*., 2012; Miyoshi and Fishell, 2011). It will be intriguing to explore how these changes in gene expression in VZ cells over time alters their neurogenic potential and cell fate capacity. In sum, this study has characterized the gene expression profile of VZ and SVZ cells in distinct neurogenic regions of the embryonic telencephalon, providing important insights into the transcriptome of distinct regional domains that may guide early neuronal fate decisions.

## METHODS

### Animals

All mouse colonies were maintained in accordance with protocols approved by the Animal Care and Use Committee at the *Eunice Kennedy Shriver* National Institute of Child Health and Human Development (NICHD). Wild-type (WT) C57BL/6 mice were obtained from The Jackson Laboratory whereas Nestin-d4-Venus (Nes-dVenus) mice were provided by RIKEN BioResource Research Center (Sunabori *et al*., 2008). For all embryonic experiments, the day on which a vaginal plug was found was considered as embryonic day (E) 0.5.

### Single-Cell Isolation

WT and Nes-dVenus pregnant dams at stage E12.5 and E14.5 were euthanized with Euthasol. Embryos were removed and incubated on ice in oxygenated artificial cerebrospinal fluid (ACSF) throughout dissection. Cortices and all three GEs were microdissected and tissue collected into labeled tubes with ACSF. Tissue was pooled from ≥ 4 embryos for each experiment and was then enzymatically dissociated with 1mg/ml of Pronase (Sigma-Aldrich #10165921001) in ACSF for 15-20 mins. Pronase solution was removed and 1-2 ml of reconstitution solution (ACSF + 1:100 fetal bovine serum (FBS) + 0.01% DNase) was added to each tube before mechanically dissociating with fire-polished glass pipettes of large, medium and small-bore openings. DAPI (1 μl) and Draq5 (5 μM, ThermoScientific #62251) were then added and cell solution was passed through a pre-wetted 35 μm filter prior to sorting (Nes-dVenus) or cell counting (WT).

### Fluorescence-activated Cell Sorting

Dissociated cell solution from Nes-dVenus mice were sorted with Beckman Coulter MoFlo Astrios cell sorter or Sony SH800S sorter with 100 μm chips. Cells were first gated with forward scatter (FSC) vs. side scatter (SSC) to remove debris, then gated with DAPI vs. Draq5 to select for live cells (Draq5+/DAPI-), then gated with GFP 488 vs. FSC to harvest GFP-expressing cells. Cells were collected into DNA LoBind microcentrifuge tubes (Eppendorf, 022431021) containing cold oxygenated ACSF supplemented with 1% FBS.

### Single-cell RNA Sequencing and Library Generation

For each experiment, ~15,000 cells from WT or sorted Nes-dVenus mice were run through the 10X Genomics Single Cell controller. Chromium Single Cell 3’ GEM (versions 2 and 3), Library and Gel Bead Kits were used according to the manufacturer’s instructions. Libraries were sequenced on Illumina HiSeq 2500 by the NICHD Molecular Genomics Core. For the E12.5 ganglionic eminences, 3 scRNAseq experiments were performed (1 WT and 2 Nes-dGFP). For the E12.5 Ctx and the E14.5 Ctx/MGE/LGE/CGE samples, 2 scRNAseq experiments were performed (1WT and 1 Nes-dGFP). After quality control analysis was used to remove low quality cells, we analyzed > 119,000 cells with a minimum of 17,500 for each brain region at E12.5 and 11,000 for each brain region at E14.5.

### Single-cell RNA Sequencing Analysis

scRNAseq data were pre-processed to count matrix with Cell Ranger (10X Genomics) with default parameters using the mm10 mouse genome as a reference. To analyze high quality cells only, we first removed cells that had unique molecular identifier (UMI) counts over 4,500 or less than 200. Cells were also filtered based on the percentage of UMI associated with mitochondrial transcripts and number of detected genes, and we removed cells >3 median absolute deviations (MADs) from the median of the total population. For cells derived from WT mice, an additional step was taken by filtering out cells based on the percentage of UMI associated with hemoglobin subunit beta (*Hbb*) transcripts (since Nes-dVenus cells were sorted, they did not require this additional filtering step). Filtration of cells based on quality control metrics, data normalization, scaling, dimensionality reduction, highly variable feature detections, population subsetting and data integration were all performed under R using Seurat (Stuart *et al*., 2019). To identify lineage inference of scRNAseq data, Slingshot was used to determine the trajectory inference on UMAP projections (Street *et al*., 2018). For the comparative analysis of VZ, SVZ and MZ cells, the effects of cell cycle heterogeneity were mitigated by calculating cell-cycle phase scores based on known canonical markers (Nestorowa et al., 2016) and the scores were regressed out during the data scaling. To distinguish cycling and postmitotic cells from *Dcx*-expressing population subsets, *Mki67*- and *Ccnd2*-expressing cells (>0.5 the log normalized count data) were positively considered SVZ cells whereas *Mki67*- and *Ccnd2*-negative cells (<0.5 the log normalized count data) were labeled as MZ cells. To combine previously reported datasets (GES103983 and GES109796) with our datasets, SCTransform normalization method was used under Seurat v3 integration workflow. For GSE109796, CGE, dMGE, and vMGE datasets from both E12.5 and E14.5 were analyzed. Cells that expressed the dorsal telencephalic cortical markers such as *Tbr1, Eomes, Neurod2* and *Neurod6* were considered contamination from the microdissected GE samples and removed prior to SCTransform normalization.

### Fluorescent *in situ* Hybridization and immunostaining

E12.5 embryonic brains were removed and drop-fixed in 4% paraformaldehyde overnight at 4°C. Fixed brains were washed in PBS, incubated in 30% sucrose overnight at 4°C and the cryopreserved. Tissues were cryosectioned at 14-16 μm in the coronal plane. RNAscope HiPlex and HiPlexUp *in situ* hybridization assays (Advanced Cell Diagnostics) were performed according to the manufacturer’s instructions. *In situ* hybridization images were taken using Zeiss AxioImager.M2 with or without ApoTome.2. Autofluorescence signals from blood vessels in embryonic brain tissues were corrected using Photoshop (Adobe) layer masking strategies. Image outlining was performed using ImageJ’s (National Institutes of Health) Canny Edge Detector and superimpose of all images was conducted with RNAscope HiPlex Image

Registration software (Advanced Cell Diagnostics). For immunofluorescence, 16 μm sections were blocked for ≥ 1 hour at RT in blocking buffer (10% Normal Donkey Serum in PBS + 0.3% Triton X-100) and incubated at 4°C overnight with primary antibodies in blocking buffer. Sections were washed 3 × 5 min at RT in PBS and incubated ≥ 1 hour at RT with fluorescent secondary antibodies and DAPI in blocking buffer. Before mounting, sections were washed again 3 × 5 min at RT in PBS and Images were captured with Axio Imager.M2 with ApoTom.2. Primary antibodies used were rabbit anti-GFP (Invitrogen A11122, 1:400), chicken anti-doublecortin (Abcam AB153668, 1:1000) and mouse anti-Nestin (Millipore MAB353, 1:100).

## ACKNOWLEDGMENTS

We would like to thank Steven L. Coon, James R. Iben and Tianwei Li at the NICHD Molecular Genomics Core for sequencing services and initial Cell Ranger analyses. We thank Dr. Dae-sung Kim and members of the Petros lab for helpful comments on the manuscript.

## AUTHOR CONTRIBUTIONS

D.R.L and T.J.P. designed the study and wrote the manuscript. D.R.L., Y.Z. and T.J.P. performed dissections and purified nuclei. D.R.L, T.J.P. and D.M. performed flow cytometry. Y.Z. prepared single cell sequencing libraries. D.R.L., C.T.R, A.M. and R.K.D. analyzed single cell data. All authors provided feedback on the manuscript.

## DECLARATION OF INTERESTS

The authors declare no competing interests.

## DATA AND SOFTWARE AVAILABILITY

The GEO accession numbers for the sequencing data reported in the paper are GSE167013.

---

**Supplementary Figure 1.**
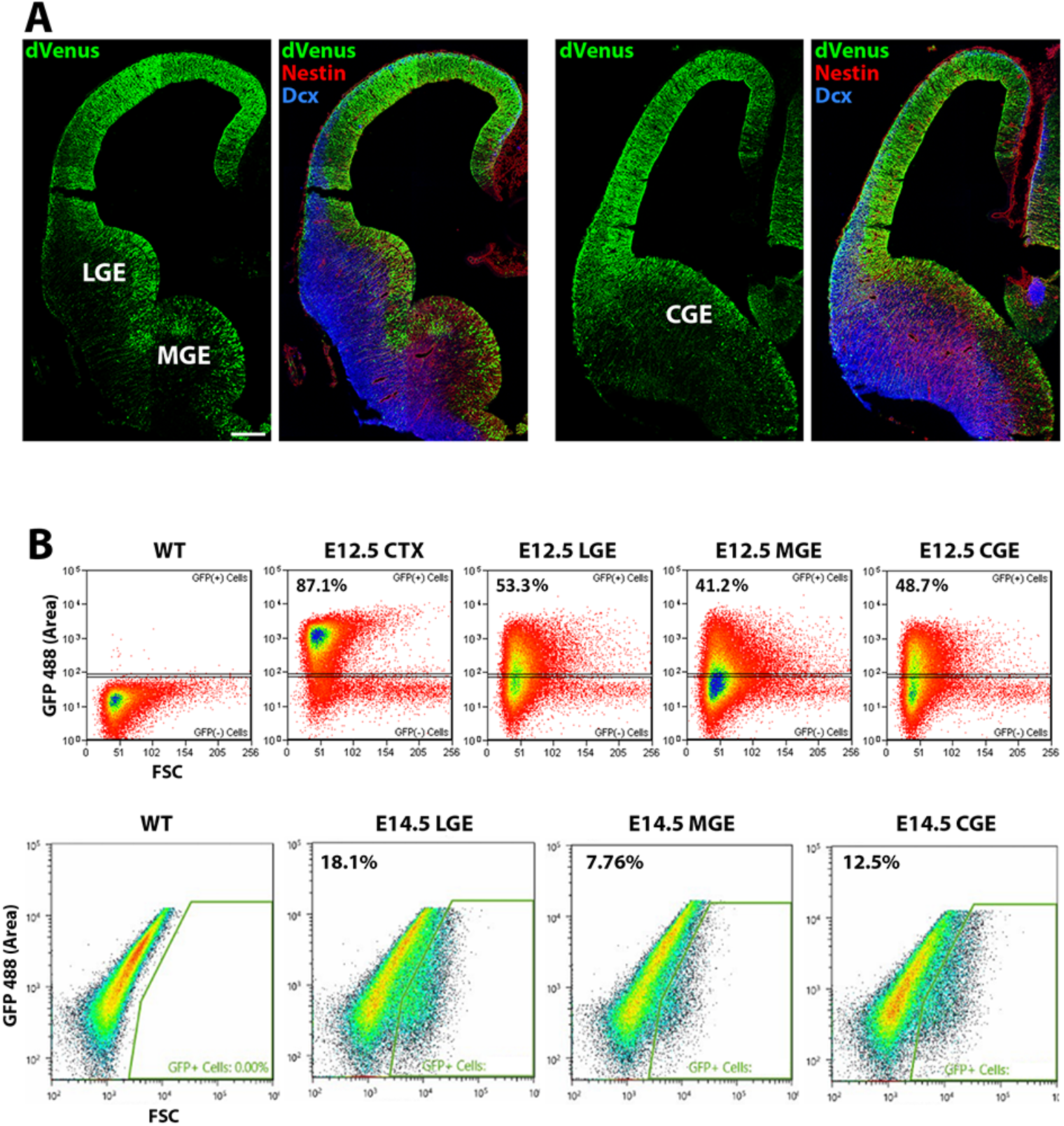
Immunostaining and FACS isolation of GFP-positive cells in Nes-dVenus mice. (A) Coronal section through E12.5 telencephalon of Nes-dVenus mouse immunostained for GFP (green), Nestin (Red) and Dcx (Blue). Scale bar = 200 μm. (B) FACS plots of depicting gating strategy to harvest GFP-positive cells from the CTX, LGE, MGE and CGE in E12.5 (top) and E14.5 (bottom) Nes-dVenus mice.

**Supplementary Figure 2.**
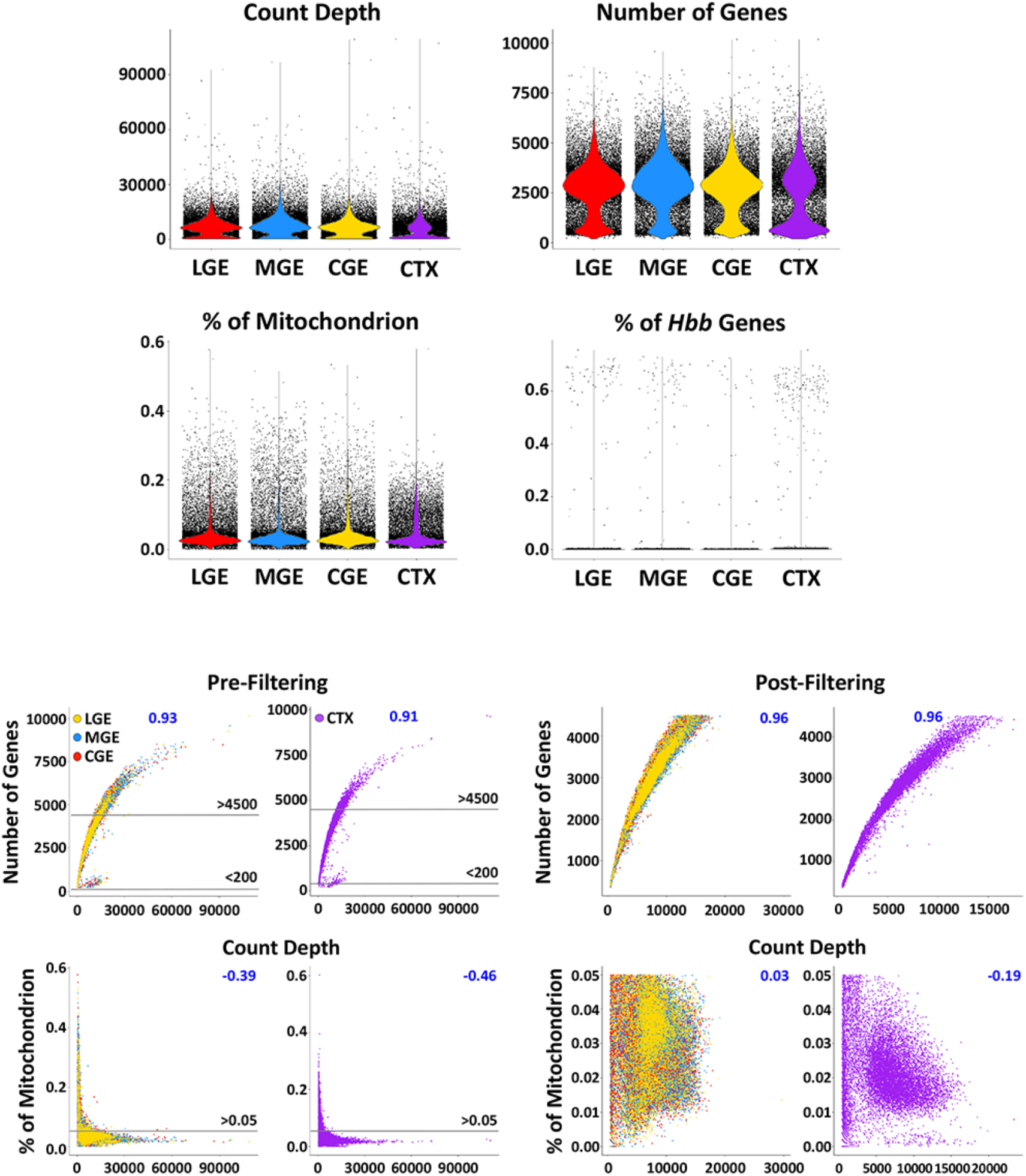
Quality-control of sequenced cells. Cells that had unique feature counts over 4500 or less than 200 were filtered out using Seurat pipeline. Mitochondrial counts over 0.05% and *Hbb* gene counts over 0.01% were also filtered out. Scatterplots showing the total number of molecules detected versus the number of genes detected before and after the filtrations (top). Additional set of scatterplots showing the total number of molecules detected versus the percentage of mitochondrial genes before and after the filtrations (bottom).

**Supplementary Figure 3.**
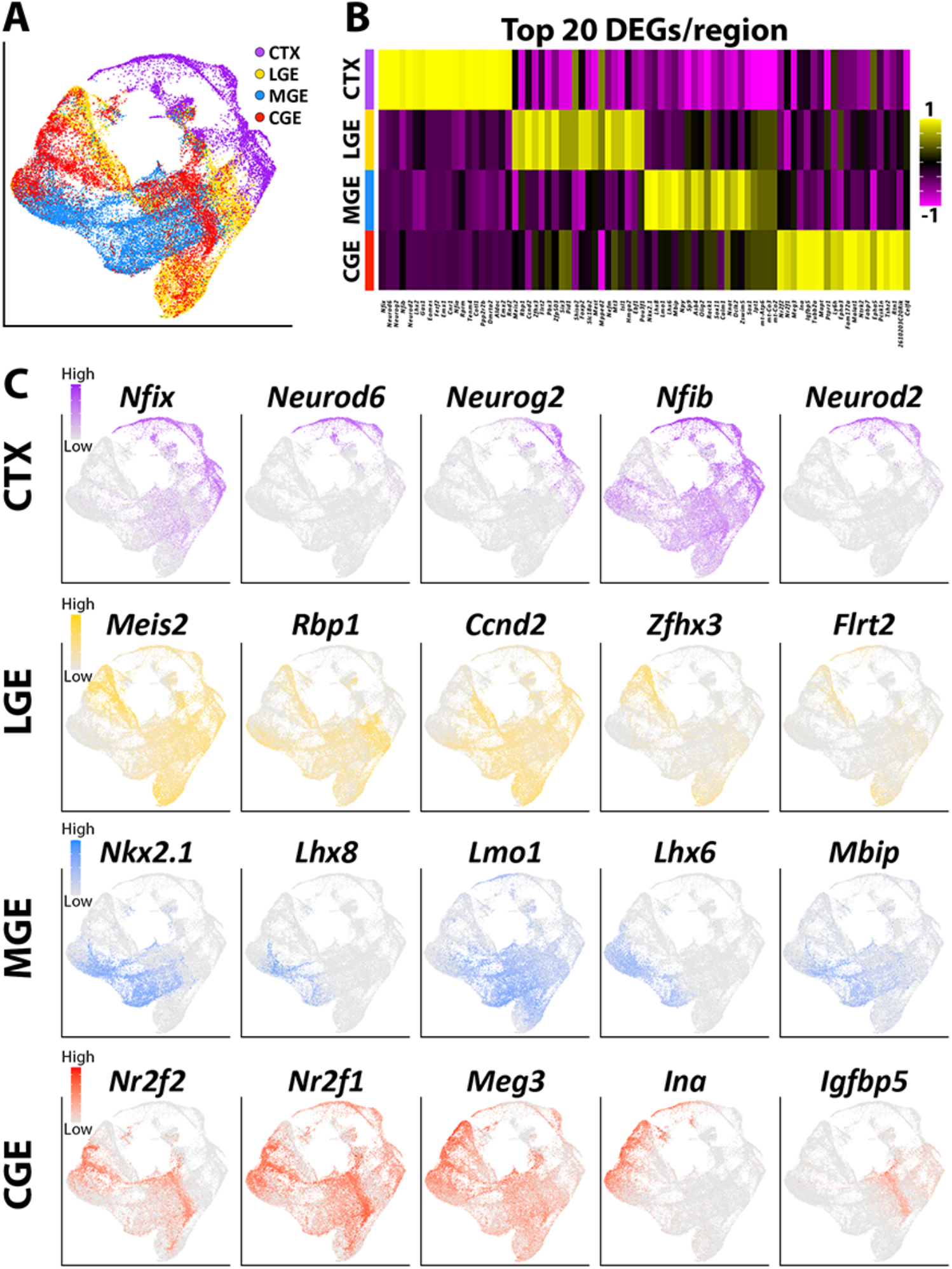
Identification of region-specific genes from each telencephalic region at E12.5. (A) UMAP plot of all cells annotated by brain region. (B) Heatmap showing expression of top 20 DEGs in each region. Each column represents averaged expression in cells, color-coded as per the color scale. (C) UMAP plots depicting the top 5 DEG expression from each region. Cells are color-coded for levels of gene expression as per the color scales.

**Supplementary Figure 4.**
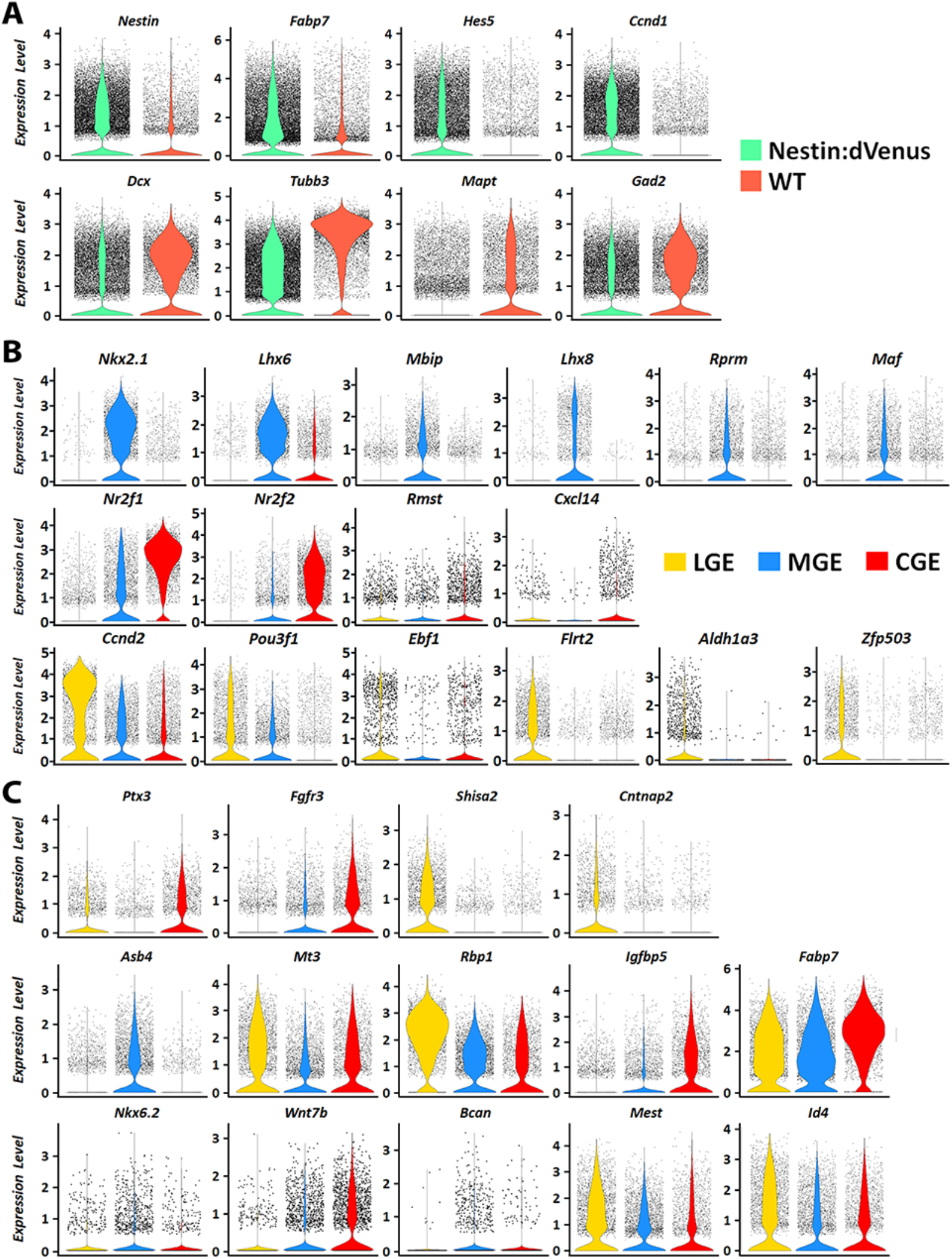
Differential gene expression between mouse lines and brain regions. (A) Violin plots showing biased expression of pan-VZ-enriched genes arising from Nes-dVenus mice and expression of SVZ/early postmitotic genes enriched in cells from WT mice. (B-C) Violin plots depicting differential gene expression profiles of cells arising from the LGE (yellow), MGE (blue) and CGE (red). Highlighted genes include those that are predominantly expressed within one GE (B) or display intriguing expression profiles based on scRNAseq and/or in situ hybridization data (C).

**Supplementary Figure 5.**
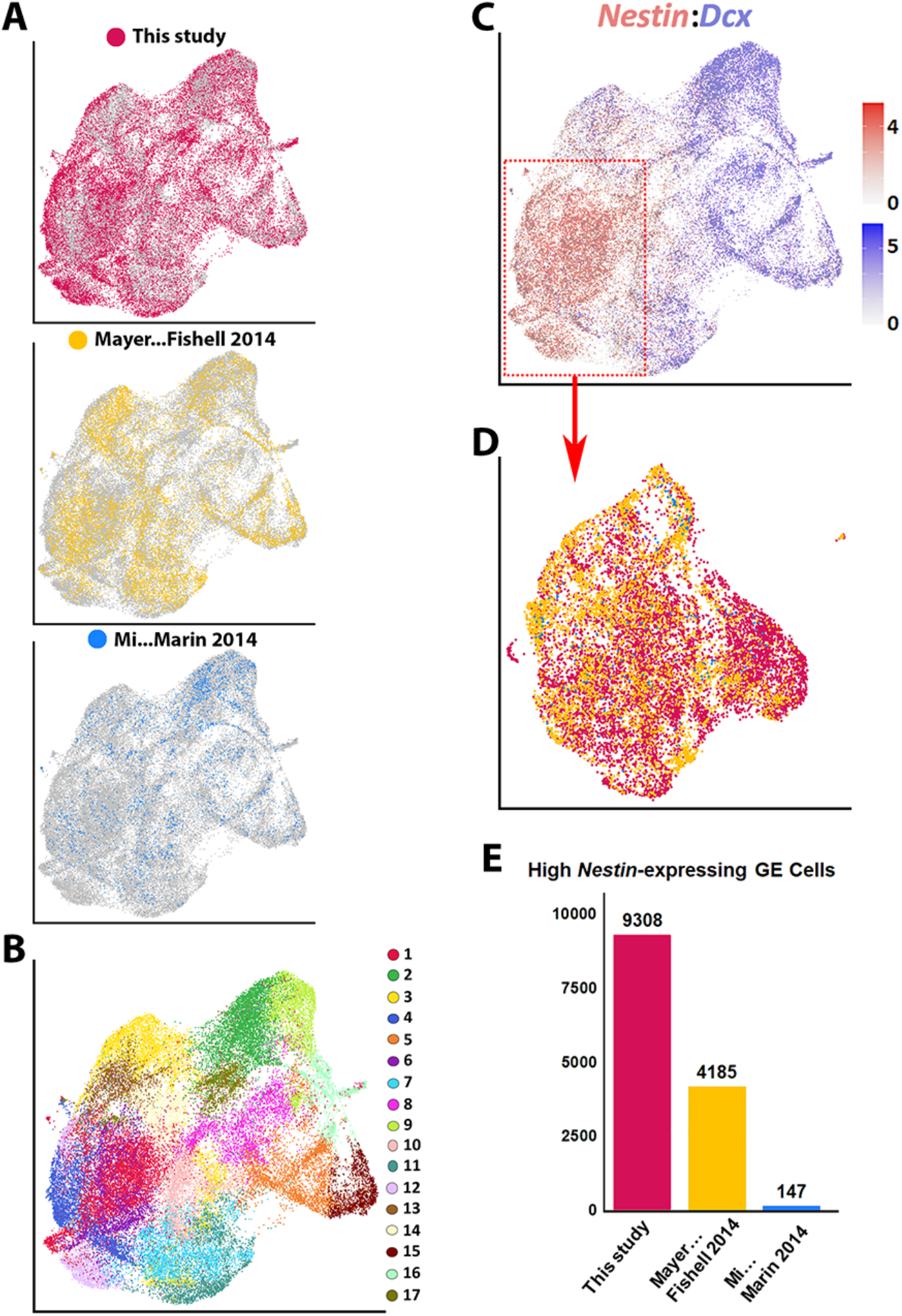
Integration and visualization of multiple embryonic mouse scRNAseq datasets. (A-C) Integration of 3 scRNAseq datasets of embryonic mouse ventral telencephalon: This study, E13.5 MGE and E14.5 LGE and CGE dataset acquired via Drop-seq from Mayer et al. 2018, and MGE and CGE dataset from two different timepoints (E12.5 and E14.5) acquired via Fluidigm C1 system by Mi et al. 2018. UMAP plots depicting all cells from the 3 studies annotated by dataset (A), putative cell clusters (B) and expression of *Nestin* and *Dcx*. (D-F) High *Nestin*-expressing cells (>1.5 normalized dataset) were extracted from the combined dataset. UMAP plot annotated by dataset (D) and bar graphs showing the total number of high *Nestin*-expressing cells present in each study (E).

**Supplementary Figure 6.**
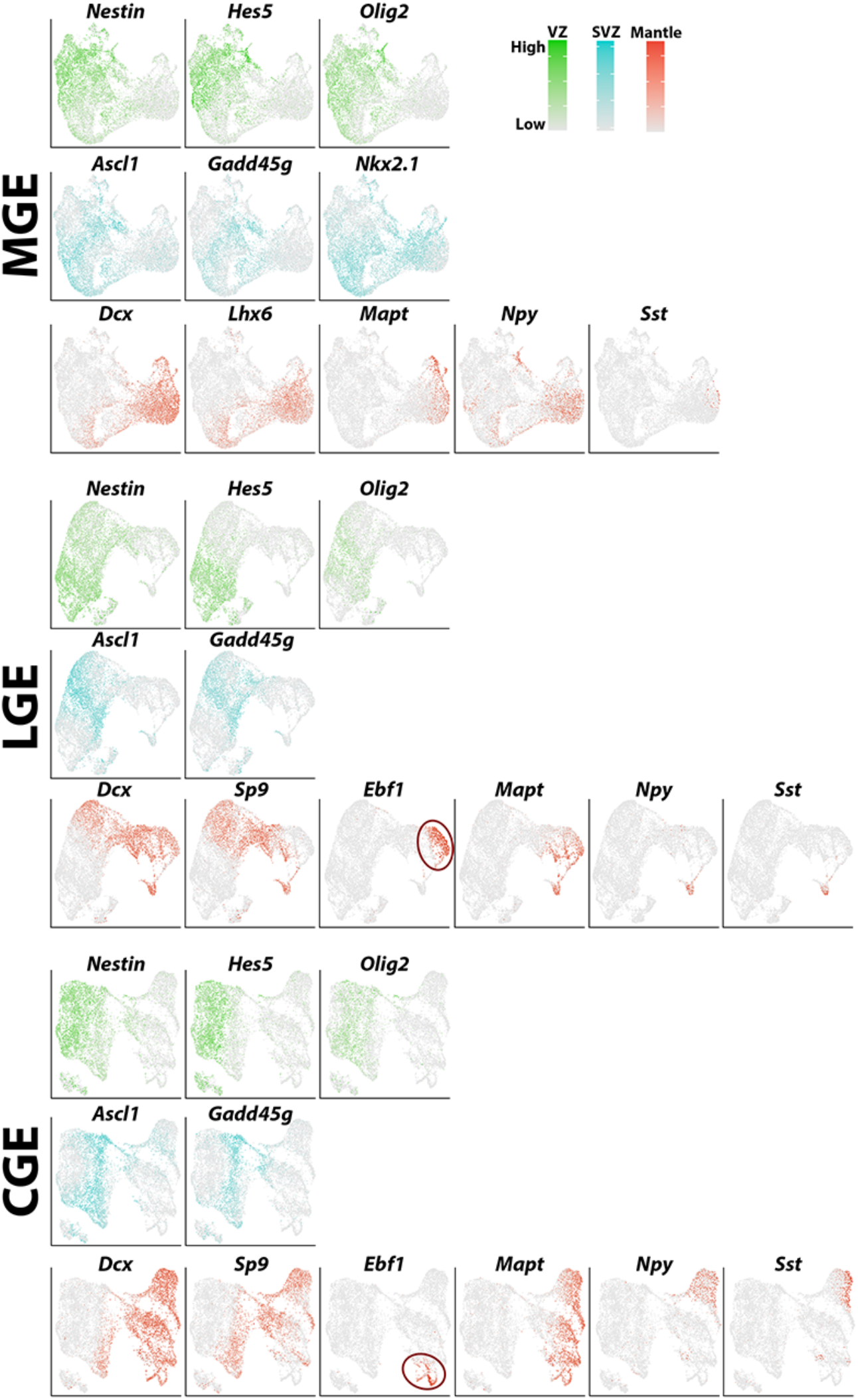
Genes defining common neurogenic cell types between brain regions. UMAP visualizations highlighting genes that help define VZ (green), SVZ (blue) and MZ (red) cell populations in the MGE, LGE and CGE. Cells are color-coded according to their levels of expression as per the color scale. Red ovals in LGE and CGE highlight the *Ebf1*-positive, *Mapt*-positive, *Sp9*-negative population of putative immature GABAergic projection neurons.

**Supplementary Figure 7.**
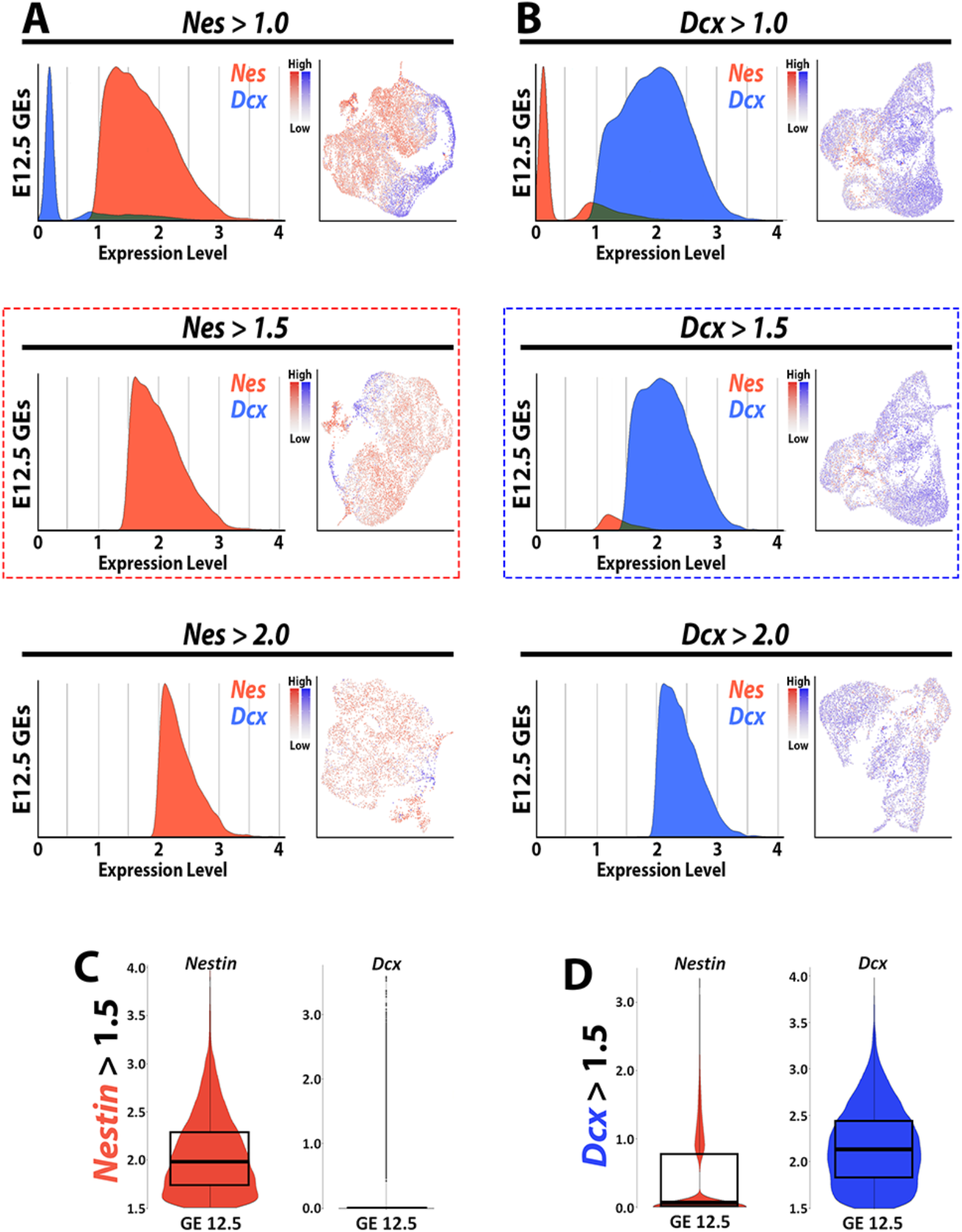
Thresholding optimization for subsetting high *Nestin*- and *Dcx*-expressing GE cells. (A-B) Ridgeline plots showing the distribution of *Nestin* (A, red) and *Dcx* (B, blue) expression when E12.5 GEs were isolated at different expression values from the log normalized count data. (C-D) Violin plot showing the median and interquartile range of *Nestin* and *Dcx* post-subsetting at *Nestin* > 1.5 (C) and *Dcx* > 1.5 (D).

**Supplementary Figure 8.**
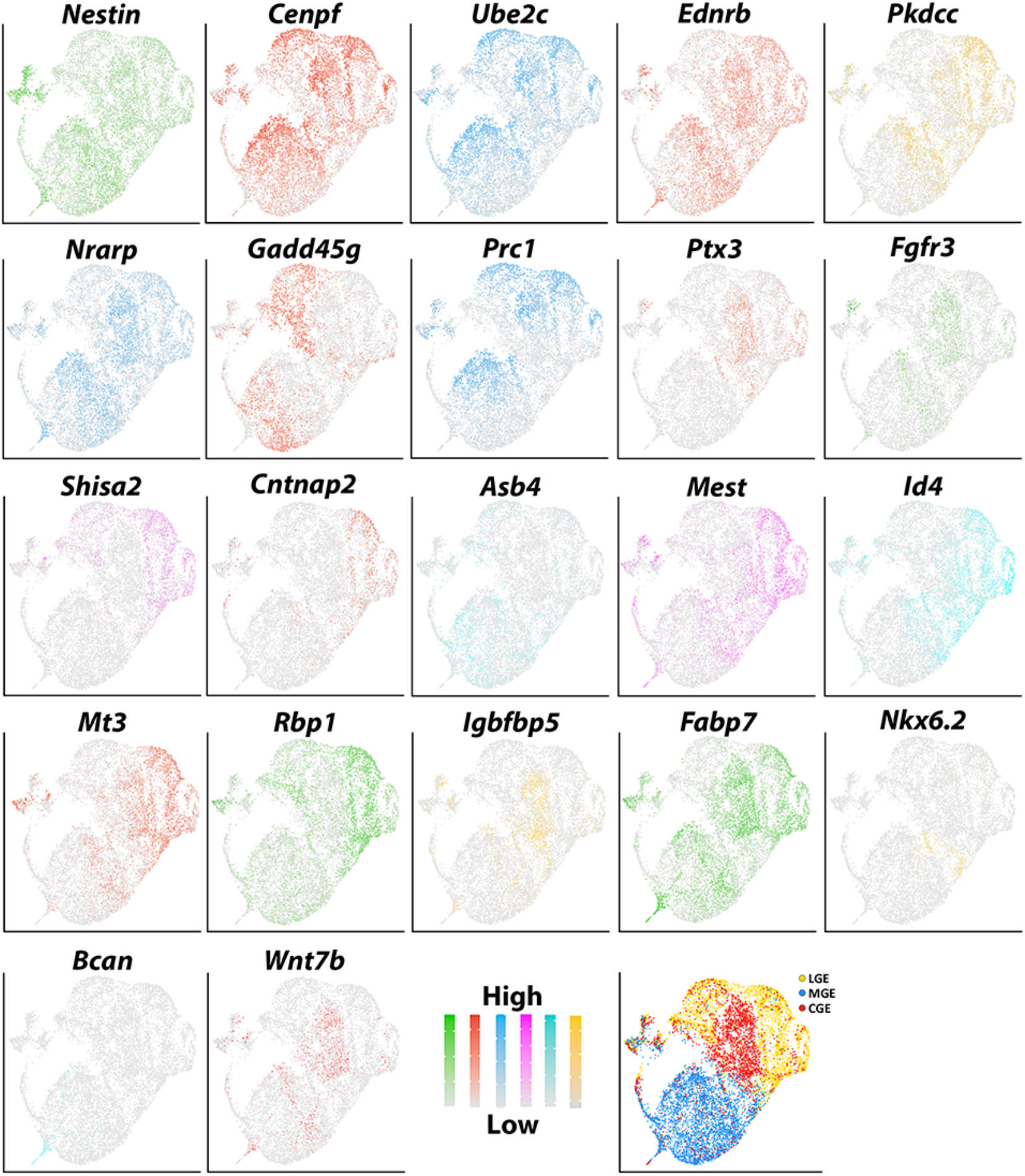
UMAP visualizations of VZ-enriched genes confirmed by in situ hybridizations. UMAP plots of VZ-enriched genes that were confirmed via RNAscope HiPlex and HiPlexUp assays, with cells annotated by brain region in last plot. Cells are color-coded for levels of gene expression as per the color scales.

**Supplementary Figure 9.**
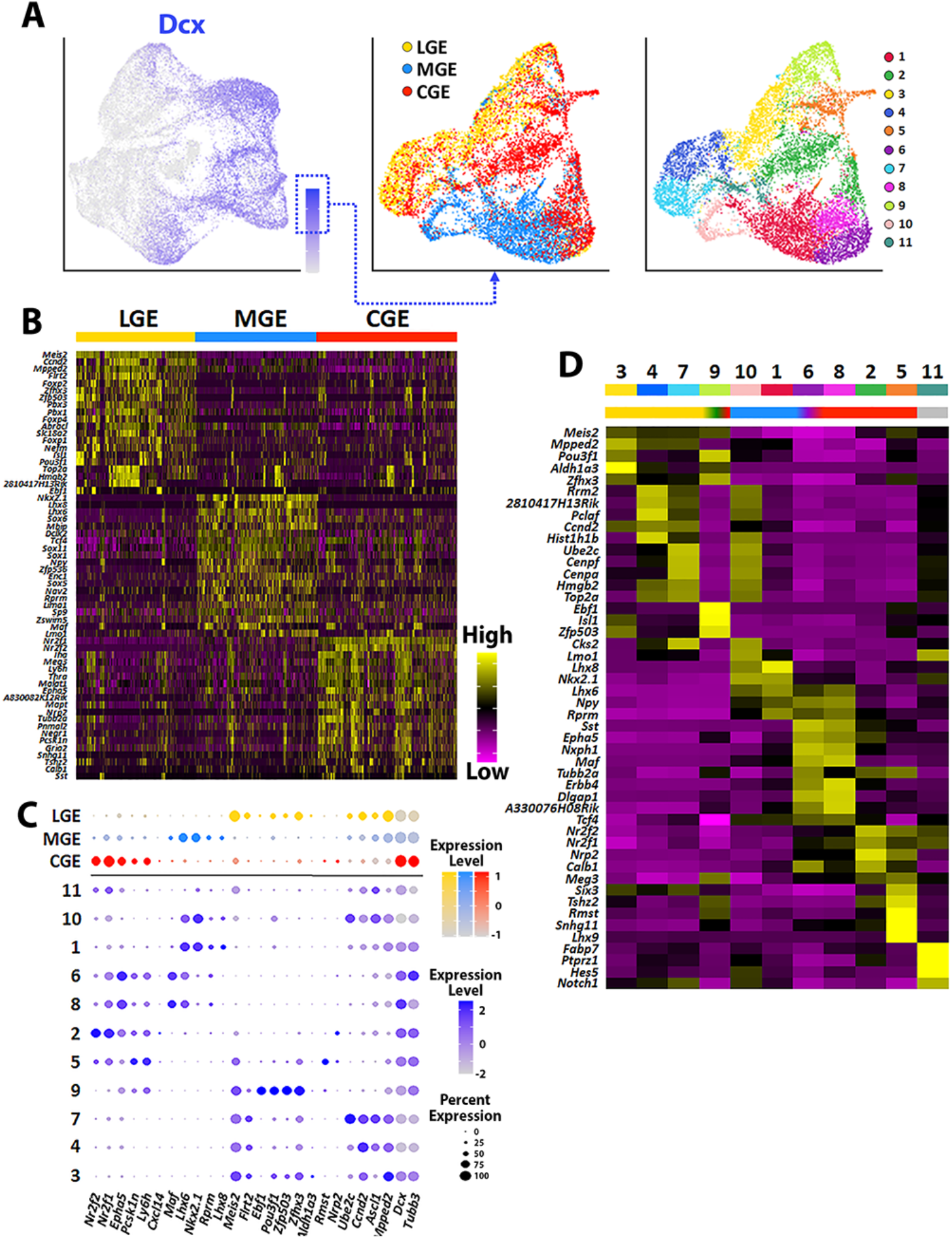
Characterization of genes enriched in high *Dcx*-expressing cells. (A) High *Dcx*-expressing cells (>1.5 normalized dataset) were extracted from the full GE dataset and replotted, annotated by brain region (middle) and putative cell clusters (right). (B) Heatmap of top 20 SVZ/MZ-enriched genes showing differential expression between LGE, MGE and CGE. Each column represents expression in a single cell, color-coded as per the color scale. (C) Dot-plot depicting expression levels of SVZ/MZ-enriched DEGs within each brain region (MGE, LGE and CGE) and cell cluster. (D) Heatmap showing expression of top 5 DEGs in each cluster from (A), with colored bar depicting whether each cluster contains cells from the LGE (yellow), MGE (blue) or CGE (red). Each column represents averaged expression in cells, color-coded as per the color scale.

**Supplementary Figure 10.**
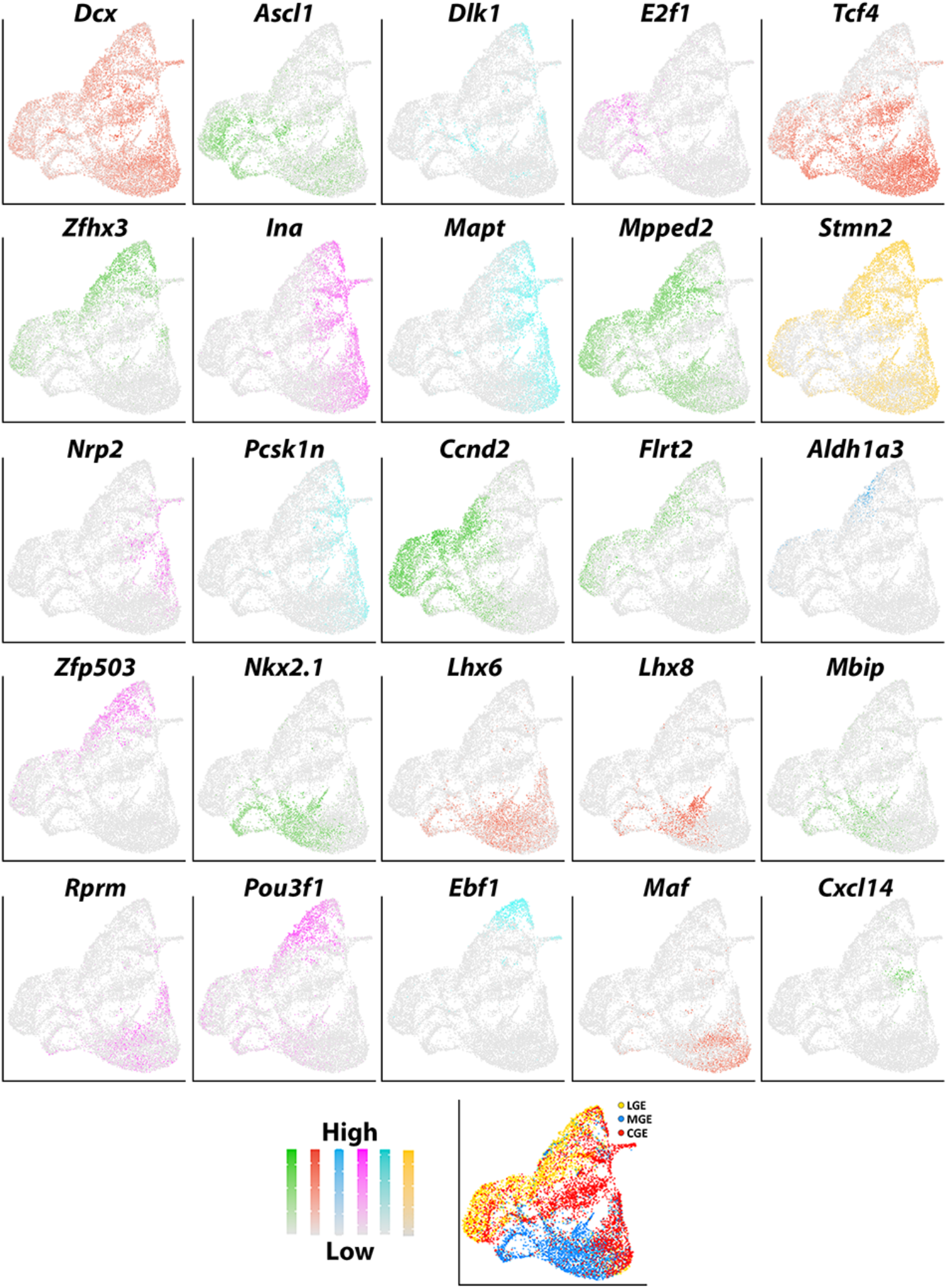
UMAP visualizations of SVZ/MZ-enriched genes confirmed by in situ hybridizations. UMAP plots of SVZ/MZ-enriched genes that were confirmed via RNAscope HiPlex and HiPlexUp assays, with cells annotated by brain region in last plot. Cells are color-coded for levels of gene expression as per the color scales.

**Supplementary Figure 11.**
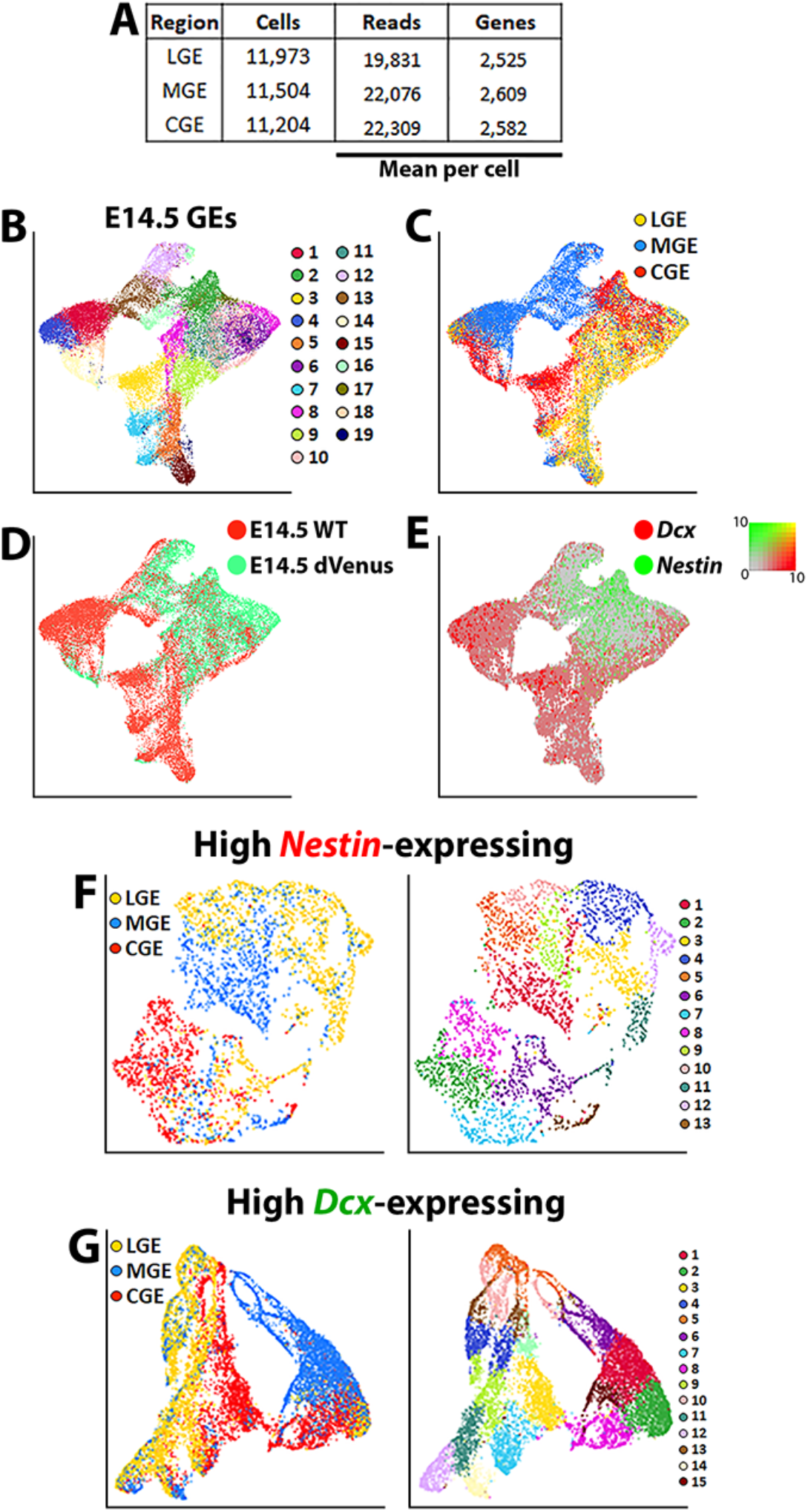
Single-cell transcriptional profiling of E14.5 ganglionic eminences. (A) Chart describing the number of cells, with mean reads per cell and genes per cell from each E14.5 ganglionic eminence. (B-E) UMAP plots of all E14.5 cells annotated by putative cell cluster (B), brain region (C), mouse line (D) and expression of *Nestin* and *Dcx*. (F) UMAP plot of high *Nestin*-expressing E14.5 cells (>1.5 normalized dataset) annotated by brain region (left) and putative cell cluster (right). (G) UMAP plot of high *Dcx*-expressing E14.5 cells (>1.5 normalized dataset) annotated by brain region (left) and putative cell cluster (right).

**Supplementary Figure 12.**
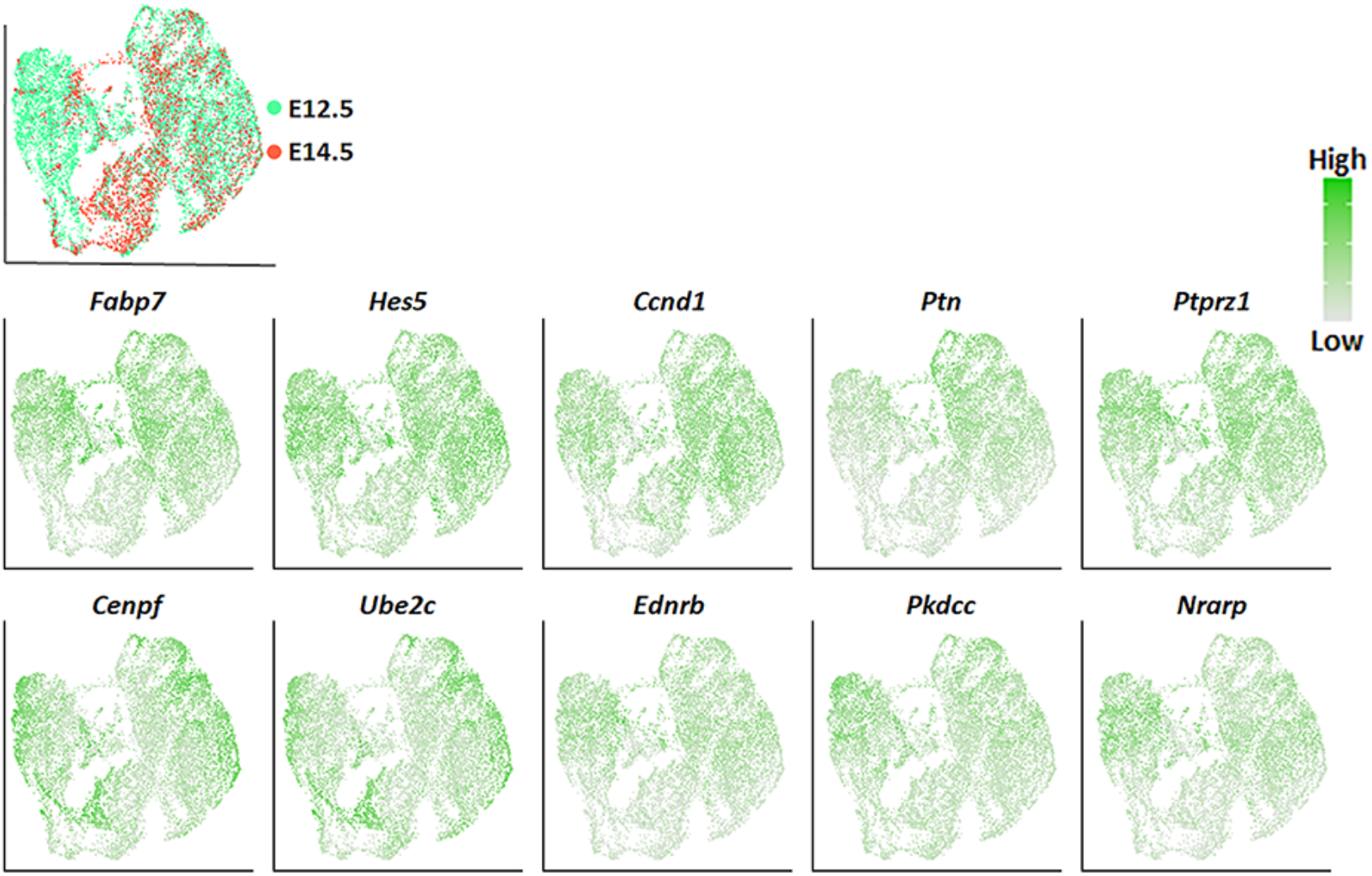
Expression of pan-VZ genes in high *Nestin*-expressing E12.5 and E14.5 cells. UMAP visualizations show a high level of pan-VZ gene expressions throughout the high *Nestin* subset E12.5 and E14.5 dataset.

## Notes

### Competing Interest Statement

The authors have declared no competing interest.

